# Non-coding RNA controls long-range enhancer-promoter interactions

**DOI:** 10.64898/2026.11.10.765568

**Authors:** Spyridoula Sagropoulou, Taras Velychko, Lara Laparra Cuervo, Simon Stitzinger, Jaschka Nicol, James L. Walshe, Yumeng Yan, Shyam Ramasamy, Lotta tom Dieck, Fenja Bahr, Olexandr Dybkov, Henning Urlaub, Kristina Žumer, Michael Lidschreiber, A. Marieke Oudelaar

**Affiliations:** Max Planck Institute of Immunobiology and Epigenetics, Department of Genome Biology, Freiburg, Germany; Georg August University of Göttingen, Göttingen, Germany; Max Planck Institute for Multidisciplinary Sciences, Department of Molecular Biology, Göttingen, Germany; Max Planck Institute for Immunobiology and Epigenetics, Department of Biological Physics, Freiburg, Germany; Max Planck Institute for Multidisciplinary Sciences, Bioanalytical Mass Spectrometry Research Group, Göttingen, Germany; University Medical Center Göttingen, Department of Pathology, Bioanalytics Group, Göttingen, Germany

## Abstract

Transcription regulation depends on interactions between enhancers and gene promoters, which often occur over large genomic distances. The molecular drivers of long-range enhancer-promoter interactions remain poorly understood. Enhancers themselves are known to be transcribed into non-coding RNAs, but the function of these transcripts is unclear. Here, we use acute genetic and chemical perturbations of transcription elongation and termination to study the role of non-coding RNA in enhancer-promoter interactions. By integrating high-resolution 3D genome mapping, transcriptomics, proteomics, biochemistry and live-cell super-resolution imaging in these perturbation contexts, we demonstrate that a strong increase in RNA molecules at enhancers disrupts their long-range interactions with gene promoters, whereas decreased RNA levels lead to stabilization of these interactions. Using separation-of-function perturbations, we show that this regulatory function of RNA is independent from the process of transcription and the proteins recruited to initiate transcription. Functionally, we show that RNA-mediated disruption of enhancer-promoter interactions is associated with reduced transcription of target genes, indicating a regulatory feedback mechanism. Together, our results uncover an additional regulatory principle of enhancer-promoter interactions mediated by non-coding RNA.

## INTRODUCTION

Enhancers play a central role in the regulation of gene expression in multicellular eukaryotes. These *cis*-regulatory elements recruit transcription factors and co-activators and thereby promote the assembly of the pre-initiation complex on gene promoters and subsequent gene transcription by RNA polymerase II (Pol II)^1^. Enhancers are often located at relatively large distances—ranging from a few to hundreds of kilobases—from the genes they control. Enhancer-mediated gene activation is therefore dependent on the formation of specific 3D interactions between enhancers and promoters in the nuclear space^2^.

Although long-range enhancer-promoter interactions have a critical role in mammalian gene regulation, the molecular mechanisms driving these interactions remain poorly understood. Loop extrusion—an active process in which cohesin molecules translocate along chromatin and extrude progressively larger loops until halted at boundary elements such as CTCF-binding sites—has a key role in large-scale organization of the genome into topologically-associating domains (TADs) and was originally proposed to be an important contributor to long-range enhancer-mediated gene activation^3,4^. Although TAD boundaries can indeed have important context-specific roles in restricting ectopic enhancer-promoter interactions^5^, recent studies have shown that the contribution of cohesin-mediated loop extrusion to the formation of enhancer-promoter interactions is often modest^6,7^, context-dependent^8^ and restricted to very long-range enhancer-promoter pairs^9–11^. In addition to loop extrusion, protein-protein interactions between transcriptional regulators bound at enhancers and promoters have been proposed to contribute to enhancer-promoter interactions^12^. It has been suggested that these interactions form in the context of transcriptional condensates, in which large amounts of transcription factors, coactivators, and Pol II are compartmentalized and concentrated^13–16^, although the precise relationship between transcriptional condensates and enhancer-promoter interactions remains unclear^17^. In line with a role for transcriptional regulators in the formation of enhancer-promoter interactions, recent studies have shown that depletion of factors such as LDB1^18^, Mediator^19^ and Pol II^20,21^ indeed weakens these interactions. However, the effects of such perturbations on enhancer-promoter interaction frequencies are generally modest. This indicates that there is redundancy between the factors involved and/or that additional determinants of enhancer-promoter interactions remain to be identified.

It is of interest that enhancers themselves are transcribed, resulting in the production of typically short and unstable enhancer RNAs (eRNAs)^22^. Enhancer transcription is particularly prominent at super-enhancers, which are clusters of closely spaced enhancer elements that recruit high levels of transcriptional regulators^23,24^. The role of these non-coding RNAs in gene regulation is controversial^25^. In particular, it is not understood if and how non-coding transcription at *cis*-regulatory elements contributes to 3D enhancer-promoter interactions. Furthermore, it is difficult to mechanistically uncouple the functional contribution of the RNA molecules themselves, the process of transcription that produces the RNA and the regulatory proteins that are recruited to *cis*-regulatory elements to initiate their transcription.

Recent insights into the role of the Integrator and Restrictor complexes in (non-coding) RNA processing and transcription termination provide elegant new strategies to investigate the putative roles of non-coding transcription and non-coding RNA in enhancer-promoter interactions. Integrator is a multi-subunit complex that was originally discovered for its role in processing small nuclear RNAs (snRNAs)^26^ but was subsequently shown to also regulate transcription at protein-coding genes and enhancers^27–29^. RNA cleavage by its catalytic subunit INTS11 and the activity of Integrator-associated protein phosphatase 2A (PP2A) oppose pause release by Cyclin-dependent kinase 9 (CDK9) and thereby promote (promoter-proximal) transcription termination, thus attenuating mRNA and eRNA production^30–33^. The Restrictor complex is composed of ZC3H4 and WDR82, which is also a subunit of the SET1/COMPASS H3K4 methyltransferase and of the PTW/PP1 phosphatase complexes^34,35^. Restrictor does not have intrinsic endonuclease activity but has also been shown to promote early termination, in particular of upstream antisense RNAs (uaRNAs), eRNAs and other unstable non-coding RNAs, and is thereby thought to restrict non-coding transcription across the genome^36–41^.

Here, we integrate complementary methods—including high-resolution 3D genome mapping, transcriptomics, proteomics, biochemistry and live-cell super-resolution imaging—to study the role of non-coding transcription and RNA in enhancer-promoter interactions. By characterizing an extensive set of (combinatorial) acute genetic and chemical perturbations—targeting Integrator, Restrictor and CDK9—that either increase or decrease non-coding RNA transcription and that allow for effective separation-of-function between the process of transcription, the proteins involved, and the produced non-coding RNA molecules, we demonstrate a key role for non-coding RNA in modulating long-range enhancer-promoter interactions in human cells.

## RESULTS

### INTS11 depletion disrupts long-range interactions between super-enhancers and their target genes

To explore if and how Integrator-dependent RNA processing affects enhancer-promoter interactions, we used an HCT116 INTS11-dTAG cell line^42^ that allows for efficient depletion of INTS11 within 90 minutes (**Extended Data Fig. 1a**). To characterize how INTS11 depletion affects RNA synthesis, we performed TT-seq experiments^43^, which confirm that rapid depletion of INTS11 has a minor effect on transcription of protein-coding genes (**Fig. 1a, Extended Data Fig. 1b,c**) but leads to increased synthesis of snRNA, uaRNA and eRNA (**Fig. 1a-b**). In addition, INTS11 depletion causes transcriptional readthrough at protein-coding genes (**Fig. 1c**), as well as extension of uaRNA and eRNA transcription beyond annotated ends (**Fig. 1d**).

**Fig. 1.**
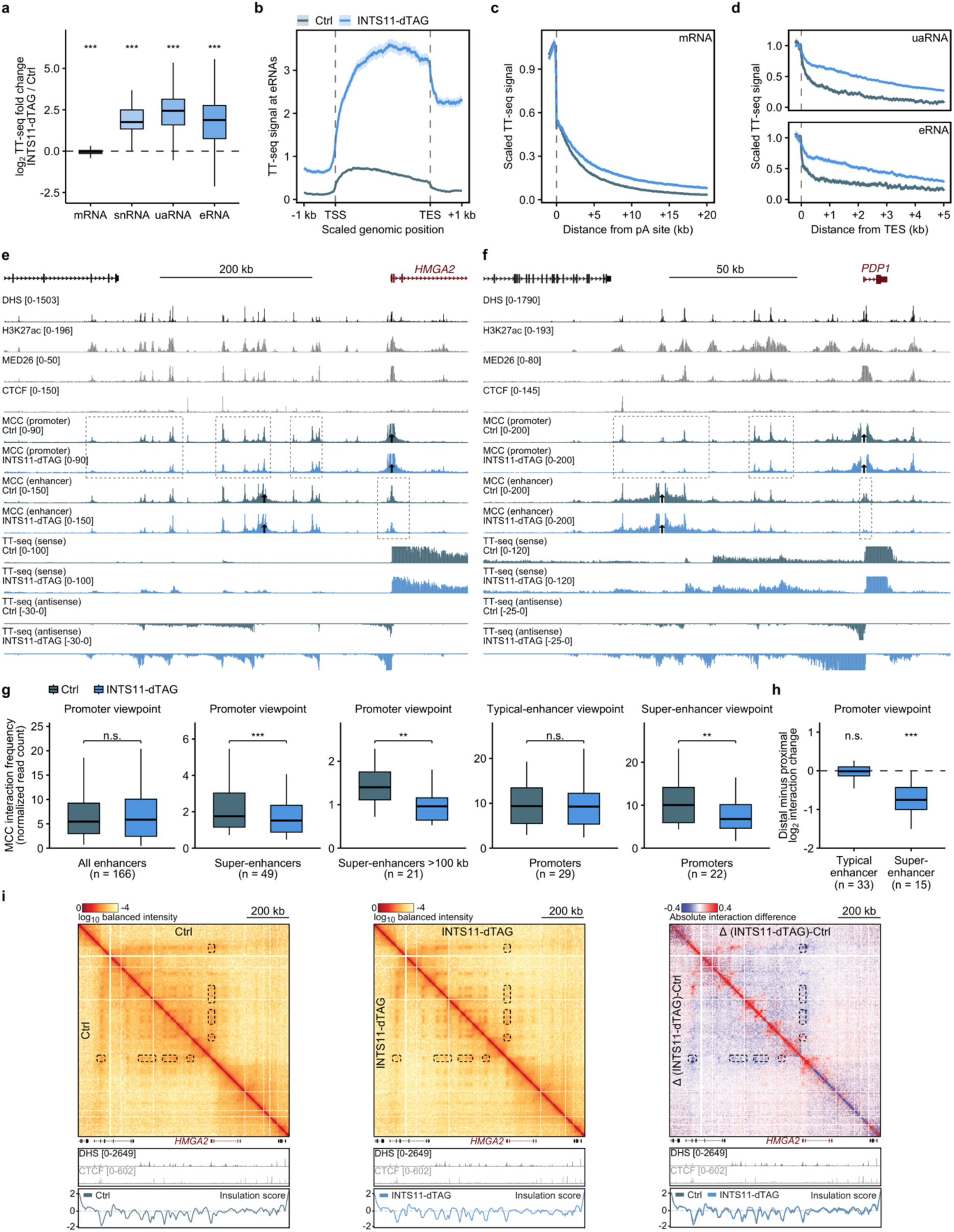
INTS11 depletion disrupts long-range interactions between super-enhancers and their target genes. **a,** TT-seq fold changes upon INTS11 depletion (INTS11-dTAG, dTAG for 90 min; Ctrl, DMSO) of mRNAs (n = 9,216), snRNAs (n = 21), uaRNAs (n = 3,591) and eRNAs (n = 3,312). **b,** TT-seq signal across eRNAs (n = 3,860), from transcription start site (TSS) to end site (TES), in Ctrl (grey) and INTS11-dTAG (blue). **c, d,** TT-seq signal downstream of the polyadenylation (pA) site of protein-coding genes (**c**; n = 7,291) and the TES of uaRNAs (d, top; n = 2,665) and eRNAs (d, bottom; n = 2,214), each condition scaled to its mean signal over the 500 bp (c) or 200 bp (d) upstream of the pA site or TES. **b-d**, Lines show 5% trimmed mean across features of merged n = 2 biological replicates and shaded areas the 95% bootstrap confidence interval. **e, f**, Micro-Capture-C (MCC) interaction profiles (mean of n = 3 biological replicates) from the viewpoint of the promoter and an enhancer (arrows) of HMGA2 (**e**; hg38 chr12:65,316,136-65,924,262) and PDP1 (**f**; chr8:93,768,209-93,950,587) in Ctrl (grey) and INTS11-dTAG (blue), with TT-seq, DNase I hypersensitive sites (DHS), H3K27ac, MED26 and CTCF; dashed boxes, regions of interest. g, MCC interaction frequencies of promoter viewpoints with all enhancers (n = 166), super-enhancers (SE; n = 49) and SE >100 kb from the promoter (n = 21), and of typical-enhancer (TE; n = 29) and SE (n = 22) viewpoints with promoters. From left, P = 0.25, 2.4 × 10^−6^, 1.5 × 10^−3^, 0.80, 1.6 × 10^−3^; n.s., not significant. h, Most distal minus most proximal interaction change (log2 INTS11-dTAG/Ctrl) per promoter with ≥2 TEs (n = 33) or SEs (n = 15). P = 0.99 (TE), 8.5 × 10^−4^ (SE). i, Tiled-MCC contact matrices of the HMGA2 locus (chr12:65,200,000-66,190,000) in Ctrl (mean of n = 3 biological replicates) and INTS11-dTAG (mean of n = 2 biological replicates) and their difference (Δ; red/blue, higher/lower in INTS11-dTAG), with DHS, CTCF and insulation scores; dashed boxes, regions of interest. Box plots: median, quartiles, 1.5 × interquartile range whiskers, no outliers. Two-sided Wilcoxon signed-rank tests (a, h, against zero; g, paired), Benjamini-Hochberg adjusted; **P < 0.01; ***P < 0.001.

To investigate how perturbation of Integrator function and the associated increase in non-coding transcription influence enhancer-promoter interactions, we performed Micro-Capture-C (MCC) experiments, which allow for generation of base-pair resolution interaction profiles for multiplexed viewpoints of interest^44,45^. Analysis of 67 interaction profiles of promoters of genes that are highly expressed in HCT116 cells shows that INTS11 depletion leads to changes in enhancer-promoter interactions. As exemplified by the *HMGA2*, *PDP1*, *MTAP* and *ITPRID2* loci, we observe that interactions between gene promoters and super-enhancers are strongly reduced upon INTS11 depletion (**Fig. 1e,f, Extended Data Fig. 1e,f**). Notably, the disruptive effect of INTS11 depletion on interactions between super-enhancers and promoters is stronger when the super-enhancers are more distal to the promoter (**Fig. 1e-g**). In contrast, interactions between gene promoters and typical enhancers are not strongly affected by INTS11 depletion (**Fig. 1g, Extended Data Fig. 1d,g,h**). These observations are confirmed by MCC interaction profiles generated from enhancer viewpoints (**Fig. 1e-g**). Quantification of the promoter-and enhancer-targeted MCC experiments shows a significant reduction in interactions between gene promoters and super-enhancers but no significant effect on interactions between gene promoters and typical enhancers (**Fig. 1g, Extended Data Fig. 1d,g,h**). Furthermore, the reduction in super-enhancer-promoter interactions is greater when the super-enhancer is located >100 kb from the gene promoter (**Fig. 1g**) and in loci containing multiple super-enhancers, interactions with the most distal super-enhancer are more affected than those with the most proximal one (**Fig. 1h**).

To better understand how 3D interactions are rewired upon INTS11 depletion, we performed Tiled-MCC (TMCC) experiments^45,46^ at loci in which we observe a reduction in enhancer-promoter interactions upon INTS11 depletion. Consistent with the MCC profiles, the TMCC matrices show a strong depletion of long-range intra-domain interactions upon INTS11 depletion, which is particularly pronounced for interactions between promoters and super-enhancers. This loss of long-range interactions is paired with a strengthening of local, short-range interactions within fine-scale compartments that align with super-enhancers (**Fig. 1i**). We conclude that INTS11 depletion leads to an increase in non-coding RNA synthesis and a decrease in long-range interactions between gene promoters and super-enhancers.

### INTS11 depletion does not reduce chromatin accessibility, recruitment of transcriptional regulators or cohesin enrichment

To disentangle whether the effects of INTS11 depletion on enhancer-promoter interactions are caused by the associated transcriptional defects or by potential accompanying reductions in chromatin accessibility or recruitment of transcriptional regulators, we characterized changes in chromatin accessibility and occupancy upon INTS11 depletion.

ATAC-seq experiments show that INTS11 depletion does not lead to decreased accessibility of gene promoters or enhancers, neither globally (**Fig. 2a-c, Extended Data Fig. 2a,b**), nor at the loci selected for MCC analysis (**Extended Data Fig. 2c)**. Rather, accessibility of *cis*-regulatory elements is overall slightly increased. This indicates that reduced enhancer-promoter interactions upon INTS11 depletion are not caused by chromatin closing or reduced accessibility for transcriptional regulators to bind to promoters and enhancers. To further confirm this, we performed ChIP analysis of MED26, a subunit of the Mediator complex, which is thought to be recruited to enhancers and promoters via transcription factors and contributes to enhancer-promoter interactions^19,47^. This analysis shows that MED26 occupancy is also not reduced upon INTS11 depletion but rather slightly increased at promoters and enhancers, both globally and at the loci selected for MCC analysis (**Fig. 2a,d,e, Extended Data Fig. 2d-f**).

**Fig. 2.**
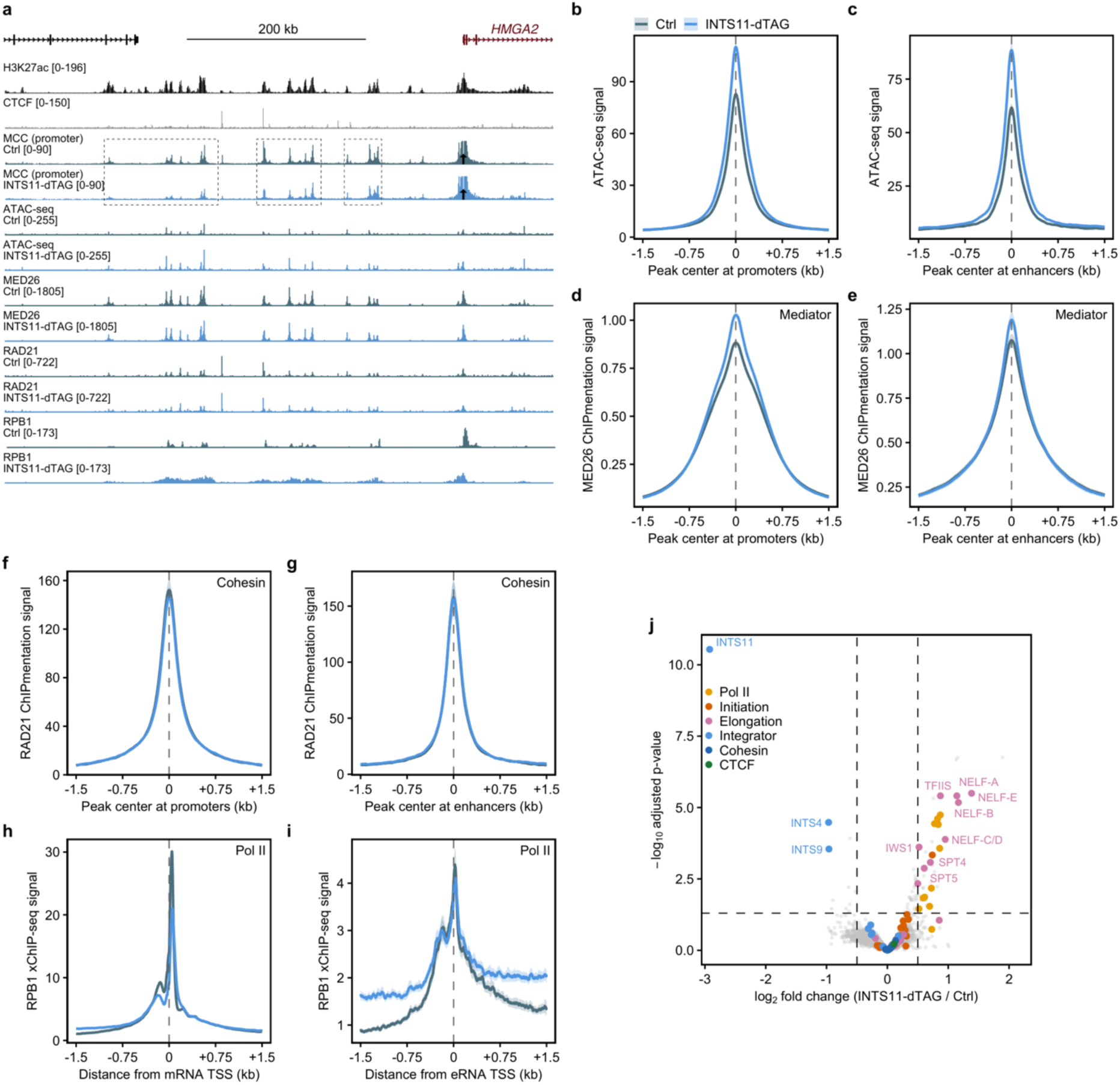
INTS11 depletion does not reduce chromatin accessibility, recruitment of transcriptional regulators or cohesin enrichment. **a,** MCC interaction profiles from the viewpoint of the HMGA2 promoter and ATAC-seq, MED26 ChIPmentation, RAD21 ChIPmentation, and RPB1 xChIP-seq signal in the control (Ctrl; grey) and upon INTS11 depletion for 90 min (INTS11-dTAG; blue). H3K27ac and CTCF profiles are shown above. Tracks as in Fig. 1e. **b-i**, Average signal in Ctrl (grey) and INTS11-dTAG (blue) of ATAC-seq at promoters (**b**; n = 8,017 peaks) and enhancers (**c**; n = 2,257), MED26 ChIPmentation at promoters (**d**; n = 8,208) and enhancers (**e**; n = 2,554), RAD21 ChIPmentation at promoters (**f**; n = 1,393) and enhancers (**g**; n = 719), and RPB1 xChIP-seq at mRNA (**h**; n = 9,283) and eRNA (**i**; n = 3,777) TSSs. Peaks are aligned at their center. Lines show the 5% trimmed mean across features (b-h) or untrimmed mean (i) of merged n = 3 biological replicates and shaded areas the 95% bootstrap confidence interval. j, Chromatin proteomics upon INTS11 depletion (4,454 proteins; n = 5 biological replicates). Dashed lines mark adjusted P = 0.05 and log2 fold changes of -0.5 and +0.5; 42 proteins were enriched on and 11 lost from chromatin. P values were calculated using two-sided moderated t-tests (limma) and adjusted using the Benjamini-Hochberg method. Subunits of the indicated complexes are colored.

It has been suggested that transcription can influence cohesin-mediated loop extrusion^20,48–51^, raising the possibility that the observed decreased long-range enhancer-promoter interactions upon INTS11 depletion are caused by perturbed loop extrusion due to increased interfering non-coding transcription at *cis*-regulatory elements. To explore this possibility, we performed ChIP analysis of the cohesin subunit RAD21. However, cohesin occupancy at promoters and enhancers is not changed upon INTS11 depletion (**Fig. 2a,f,g, Extended Data Fig. 2g-i**), indicating that changes in loop extrusion are unlikely to underlie the observed changes in enhancer-promoter interactions.

Pol II itself has also been implicated in the formation of enhancer-promoter interactions^20,21,52^. ChIP analysis of the RPB1 subunit of Pol II shows a decrease in Pol II binding at gene promoters upon INTS11 depletion. In contrast, Pol II at enhancer transcription start sites (TSSs) is not strongly affected and occupancy in the flanking regions is increased (**Fig. 2a,h,i, Extended Data Fig. 2j-m**), in line with increased and extended eRNA transcription upon INTS11 depletion (**Fig. 1b,d**). This indicates that—in addition to being caused by the transcriptional defects upon INTS11 depletion—the reduced enhancer-promoter interactions could in principle also be related to slightly reduced Pol II binding at the gene promoters.

The ChIP analyses are further supported by chromatin proteomics experiments (**Fig. 2j, Extended Data Fig. 2n**), which show that the only Pol II-associated proteins with reduced chromatin binding are INTS11, INTS4 and INTS9, which form the cleavage module of the Integrator complex^53^. Notably, this indicates that the changes in enhancer-promoter interactions are unlikely to be related to a direct bridging function of the Integrator complex, since binding of the remaining Integrator subunits to chromatin is not affected upon INTS11 depletion (**Extended Data Fig. 2o**). Furthermore, the proteomics data do not show clear changes in chromatin binding of transcription factors, and no reduction of Mediator subunits, cohesin subunits or CTCF. Despite the reduction in Pol II ChIP signal at gene promoters, the proteomics data show an overall increase in Pol II and transcription elongation factors—such as negative elongation factor (NELF), DRB sensitivity-inducing factor (DSIF), SPT6, and TFIIS— on chromatin, in line with increased non-coding transcription and transcription readthrough upon INTS11 depletion (**Fig. 1a-d**).

Taken together, we conclude that the effects of INTS11 depletion on enhancer-promoter interactions are not caused by accompanying losses in chromatin accessibility, recruitment of Mediator, interference of cohesin-mediated loop extrusion or a potential direct bridging function of the Integrator complex. Possible remaining explanations are: (1) increased transcription of promoter-proximal and enhancer regions: a higher number of transcribing polymerases ploughing through these regions may physically interfere with the molecular interactions underlying enhancer-promoter proximity; (2) increased non-coding RNA molecules bound to promoter-proximal and enhancer chromatin: RNA molecules may sterically or electrostatically disrupt molecular interactions that contribute to enhancer-promoter proximity; (3) decreased Pol II at gene promoters: this may impair direct molecular bridging by Pol II and lead to decreased enhancer-promoter proximity. Since INTS11 depletion affects all three potential mechanisms simultaneously, their individual contributions cannot be distinguished by studying this perturbation in isolation.

### Decoupling transcription, Pol II binding and RNA levels

To identify the molecular mechanism by which INTS11 depletion impairs enhancer-promoter interactions, we sought a way to decouple the effects on transcription, Pol II binding and RNA levels that are induced by INTS11 depletion. To this end, we combined 90 min INTS11 depletion with treatment with the CDK9 inhibitor NVP2^54^ during the final 45 min. CDK9—the catalytic subunit of Positive Transcription Elongation Factor b (P-TEFb)—phosphorylates the Pol II C-terminal domain (CTD), DSIF and NELF to release promoter-proximally paused Pol II molecules into productive elongation^55,56^. CDK9 inhibition therefore suppresses the entry to productive elongation, which is expected to counteract the increase in non-coding transcription in INTS11-depleted cells.

To validate effective CDK9 inhibition, we performed TT-seq experiments. As expected, these experiments show a strong decrease in uaRNA and eRNA synthesis for combined INTS11 depletion and CDK9 inhibition compared to INTS11 depletion alone (**Fig. 3a,b**). Of note, mRNA synthesis levels, which are not strongly affected by INTS11 depletion (**Fig. 1a**), are strongly decreased when INTS11 depletion is combined with CDK9 inhibition (**Fig. 3a, Extended Data Fig. 3a**). Chromatin accessibility (**Extended Data Fig. 3b-f**) and cohesin occupancy (**Extended Data Fig. 3g-k**) are not reduced upon combined INTS11 depletion and CDK9 inhibition. Chromatin proteomics experiments show the same selective loss of the Integrator cleavage module upon combined INTS11 depletion and CDK9 inhibition as was observed for INTS11 depletion alone, together with further accumulation of NELF on chromatin and modest loss of elongation factors from chromatin (**Extended Data Fig. 3l,m**). However, notably, Pol II occupancy is increased downstream of promoter-proximal pause positions and at enhancer regions, and is largely restored at the super-enhancer-associated gene promoters analyzed by MCC (**Fig. 3c,d, Extended Data Fig. 3n-p**), as exemplified by the *HMGA2* and *PDP1* loci (**Fig. 3e,f**). Together, this indicates that upon CDK9 inhibition in the context of INTS11 depletion, more Pol II molecules are transcriptionally engaged, while producing less RNA, consistent with impaired release into productive elongation.

**Fig. 3.**
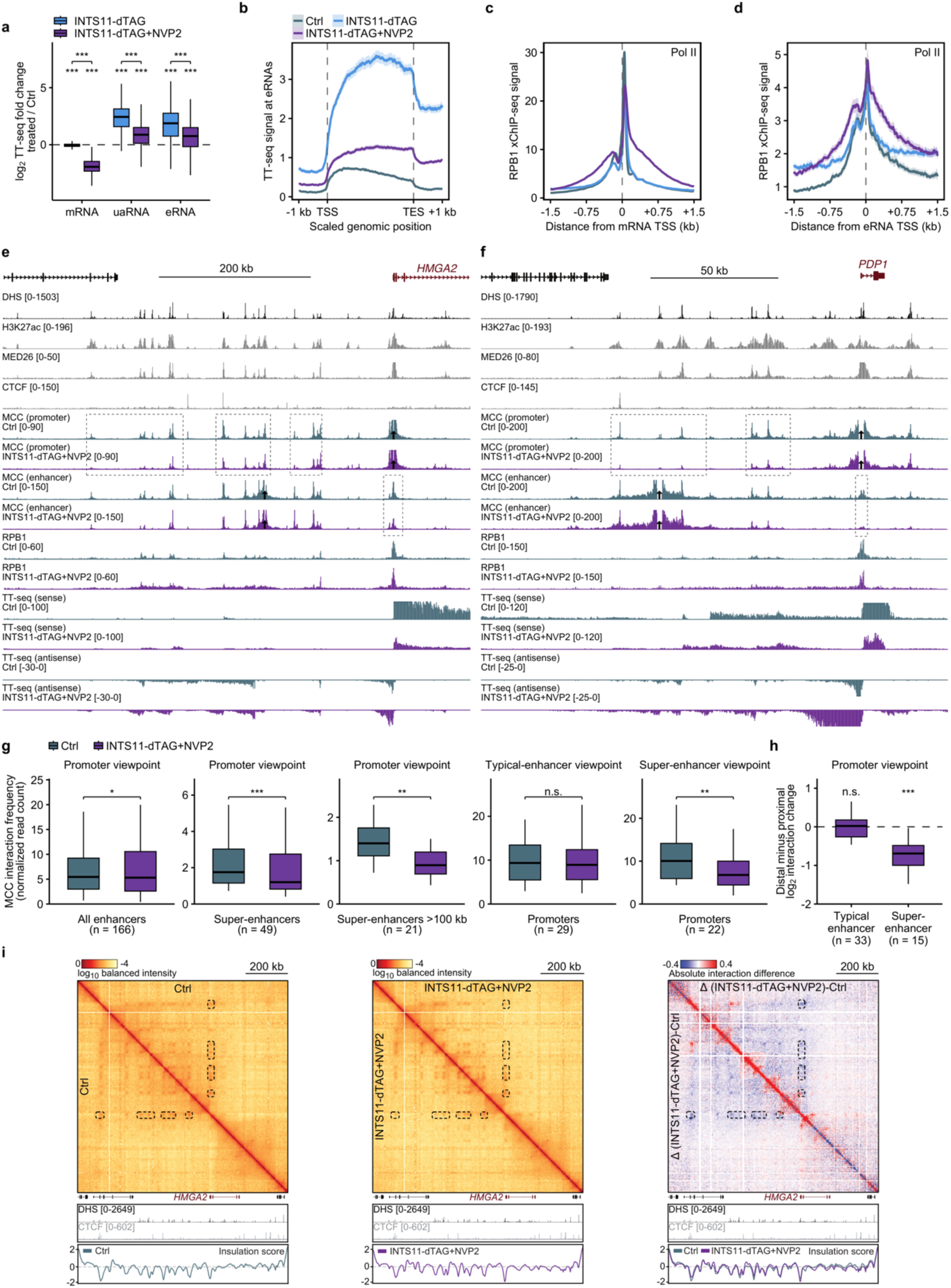
Reducing non-coding transcription does not restore enhancer-promoter interactions upon INTS11 depletion. **a,** TT-seq fold changes of mRNAs (n = 9,216), uaRNAs (n = 3,591) and eRNAs (n = 3,312) after INTS11 depletion for 90 min (INTS11-dTAG; blue) or INTS11 depletion with the CDK9 inhibitor NVP2 for the final 45 min (INTS11-dTAG+NVP2; purple). Box plots as in Fig. 1. Two-sided Wilcoxon signed-rank tests against zero (stars above boxes) and between treatments (brackets), Benjamini-Hochberg adjusted. **b,** TT-seq signal across eRNAs (n = 3,860); profiles as in Fig. 1b. **c, d**, RPB1 xChIP-seq signal at mRNA (**c**; n = 9,283) and eRNA (**d**; n = 3,777) TSSs; profiles as in Fig. 2. **e,** MCC interaction profiles (mean of n = 3 biological replicates), from the HMGA2 promoter and enhancer viewpoints, with RPB1 xChIP-seq and TT-seq signal, in Ctrl (grey) and INTS11-dTAG+NVP2 (purple). Tracks as in Fig. 1e. **f,** As in **e,** for the PDP1 locus. Tracks as in Fig. 1f. **g,** MCC interaction frequencies as in Fig. 1g, in Ctrl and INTS11-dTAG+NVP2. From left, P = 0.017, 5.1 × 10^−6^, 2.9 × 10^−3^, 0.93, and 4.8 × 10^−3^. Note that there is a very slight increase in promoter interactions with all enhancers but a strong decrease in promoter interactions with super-enhancers. **h,** Within-locus interaction changes, as in Fig. 1h, in INTS11-dTAG+NVP2. P = 0.71 (TE) and 1.2 × 10^−4^ (SE). i, Tiled-MCC contact matrices of the HMGA2 locus in Ctrl (mean of n = 3 biological replicates) and INTS11-dTAG+NVP2 (mean of n = 2 biological replicates) and the differential matrix (Δ (INTS11-dTAG+NVP2)−Ctrl), as in Fig. 1i. n.s., not significant; *P < 0.05; **P < 0.01; ***P < 0.001.

To investigate whether INTS11 depletion still causes reduced interactions between promoters and super-enhancers under conditions in which there is less increased transcription of these *cis*-regulatory elements and Pol II binding is largely restored, we performed MCC experiments upon combined INTS11 depletion and CDK9 inhibition, targeting the same set of promoters and enhancers. Notably, we observe an almost identical effect on enhancer-promoter interactions as following INTS11 depletion in isolation, with a strong reduction in interactions between super-enhancers and promoters, which is augmented when the super-enhancer is located more distal to the promoter, but no reduction in interactions between promoters and typical enhancers (**Fig. 3e-i**).

Taken together, our experiments show that upon combined INTS11 depletion and CDK9 inhibition, non-coding transcription is less strongly increased and Pol II binding is largely restored, yet the effects on enhancer-promoter interactions are almost identical to depletion of INTS11 alone. This suggests that the changes in enhancer-promoter interactions upon INTS11 depletion are not due to a perturbing effect of increased transcription through *cis*-regulatory elements or reduced Pol II occupancy at these elements.

### Increased chromatin-associated RNA levels upon INTS11 depletion lead to reduced enhancer-promoter interactions and enhancer-mediated gene activation

TT-seq measures RNA synthesized during a short period of 4sU labeling and therefore informs on RNA synthesis (i.e., transcription) rates^43^, which are strongly reduced upon CDK9 inhibition (**Fig. 3a**). It is important to note that TT-seq signal depends not only on the number of transcribing polymerases but also on their elongation velocity^57^ which is reduced upon combined INTS11 depletion and CDK9 inhibition (**Extended Data Fig. 4a**). A decrease in RNA synthesis therefore does not necessarily imply a decrease in the amount of RNA present on chromatin. To identify potential accumulation of non-coding RNA on chromatin across the tested conditions, we performed chromatin-associated RNA-sequencing (chrRNA-seq) experiments. Notably, these experiments show that chromatin-associated non-coding RNA levels are strongly increased upon both INTS11 depletion alone and upon INTS11 depletion in combination with CDK9 inhibition (**Fig. 4a,b**). In addition, direct comparison of TT-seq and chrRNA-seq signal across the genome between these two conditions shows that non-coding RNA transcription is much lower upon combined INTS11 depletion and CDK9 inhibition compared to INTS11 depletion alone, whereas chromatin-associated non-coding RNA levels are very similar in both conditions (**Extended Data Fig. 4b,c**). While these data strongly suggest that increased chromatin-associated non-coding RNA underlies the reduced enhancer-promoter interactions observed in both these conditions, we cannot fully rule out a perturbing effect of actively transcribing polymerases, since non-coding RNA transcription remains approximately 1.7-fold above the DMSO control under combined INTS11 depletion and CDK9 inhibition (**Fig. 3a,b**).

**Fig. 4.**
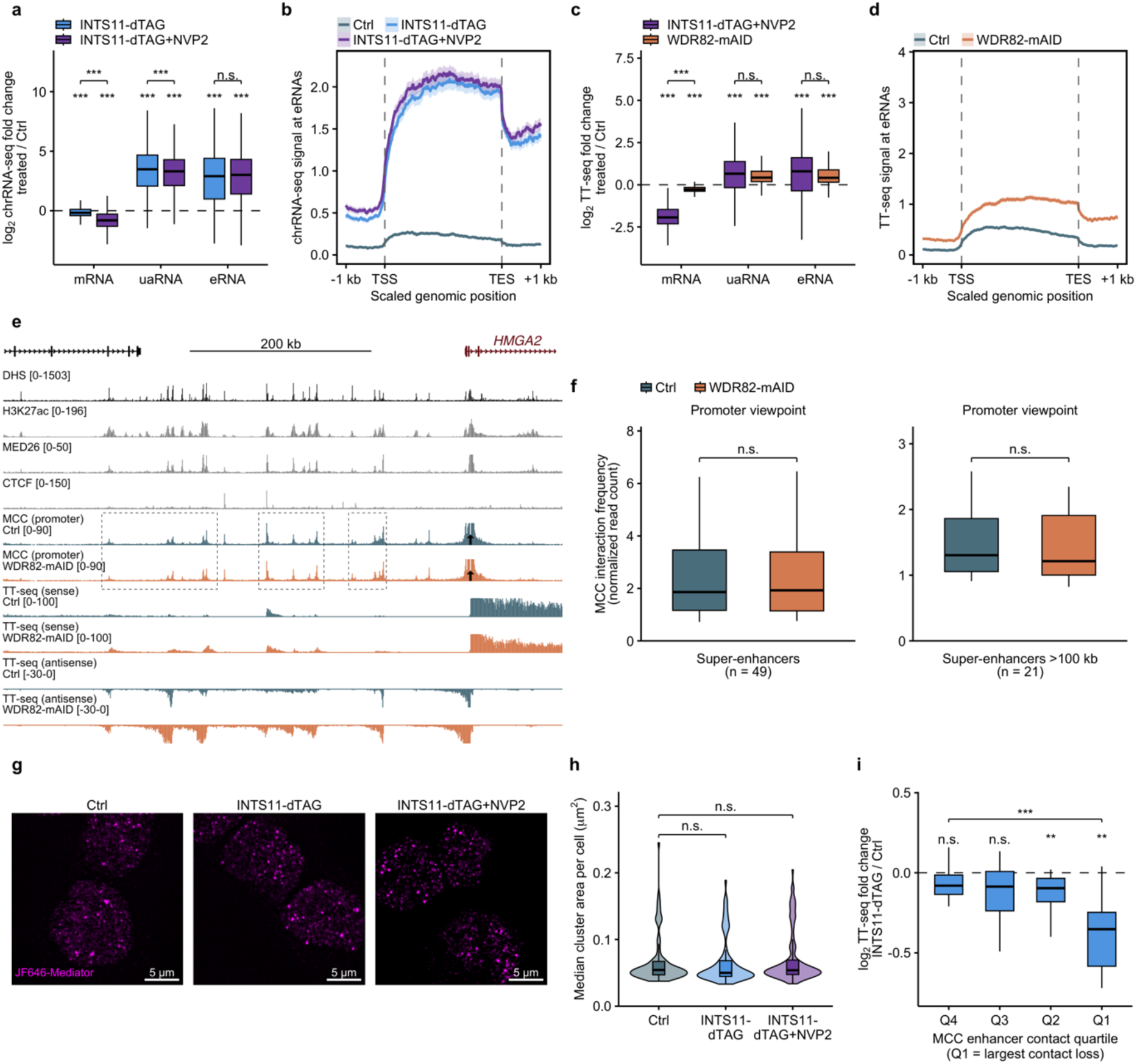
Increased chromatin-associated RNA levels upon INTS11 depletion lead to reduced enhancer-promoter interactions and enhancer-mediated gene activation. **a,** chrRNA-seq fold changes of mRNAs (n = 9,213), uaRNAs (n = 3,539) and eRNAs (n = 3,182) in INTS11-dTAG (blue) or INTS11-dTAG+NVP2 (purple). Box plots and tests as in Fig. 3a. **b,** chrRNA-seq signal across eRNAs (n = 3,860). Profiles as in Fig. 1b. c, TT-seq fold changes in INTS11-dTAG+NVP2 (purple) or in WDR82-mAID (orange), each against its own control, of mRNAs (n = 9,189), uaRNAs (n = 2,268) and eRNAs (n = 1,831) detected in both conditions. Box plots and tests as in Fig. 3a. **d,** As **b,** for TT-seq in Ctrl and WDR82-mAID. Profiles as in Fig. 1b. e, MCC interaction profiles (mean of n = 3 biological replicates) from the HMGA2 promoter viewpoint and TT-seq in Ctrl (grey) and WDR82-mAID (orange). Tracks as in Fig. 1e. **f,** MCC interaction frequencies of promoter viewpoints with super-enhancers (SE; n = 49) and SE >100 kb away (n = 21), as in Fig. 1g. P = 0.33, 0.19. **g,** Structured illumination microscopy (SIM) of MED19 (Mediator) in INTS11-dTAG-MED19-Halo cells; scale bars, 5 µm. **h,** Median MED19 cluster area per cell in Ctrl, INTS11-dTAG and INTS11-dTAG+NVP2 (n = 94, 92 and 109 cells; clusters of 30-500 pixels, 0.029-0.48 µm²); violins with box plots. Two-sided Wilcoxon rank-sum tests against Ctrl, Benjamini-Hochberg adjusted; P = 0.43 for both. i, TT-seq fold changes in INTS11-dTAG of the MCC-targeted genes by quartile (n = 15-16) from least (Q4) to most (Q1) reduced enhancer-promoter interaction. Spearman ρ = 0.46. Two-sided Wilcoxon signed-rank tests against zero and rank-sum between quartiles (bracket, Q4 vs Q1), Benjamini-Hochberg adjusted. n.s., not significant; **P < 0.01; ***P < 0.001.

Therefore, to further disentangle the role of the active process of transcription and chromatin-associated RNA accumulation, we characterized an orthogonal perturbation that increases non-coding transcription as measured by TT-seq but produces less accumulation of chromatin-associated RNA. To this end, we used a HCT116 WDR82-mAID cell line^38^ that allows for depletion of WDR82, a subunit of the Restrictor complex, within 4 hours (**Extended Data Fig. 4d**). While WDR82 depletion mildly reduces global mRNA transcription (**Fig. 4c**), it increases non-coding RNA transcription to similar levels as observed upon combined INTS11 depletion and CDK9 inhibition (**Fig. 4c,d, Extended Data Fig. 4e-g**), including at super-enhancers (**Extended Data Fig. 4h**). In contrast, the increase in chromatin-associated non-coding RNA levels upon WDR82 depletion is much smaller compared to the changes observed in the context of INTS11 depletion, both with and without simultaneous CDK9 inhibition (**Extended Data Fig. 4e-h**).

To test whether the moderate increase in non-coding RNA transcription in the context of WDR82 depletion is associated with changes in enhancer-promoter interactions, we performed MCC experiments, targeting the same set of promoters and enhancers as in our previous experiments. Interestingly, the MCC data do not show clear changes in interactions between promoters and super-enhancers or typical enhancers upon WDR82 depletion (**Fig. 4e,f**). Thus, WDR82 depletion and combined INTS11 depletion and CDK9 inhibition increase non-coding RNA transcription to a similar extent (**Fig. 4c**), but only the latter is accompanied by a strong increase in chromatin-associated non-coding RNA (**Fig. 4a, Extended Data Fig. 4e-h**) and by loss of enhancer-promoter interactions (**Fig. 3g**). We therefore conclude that the reduction in enhancer-promoter interactions that we observe upon INTS11 depletion is caused by the accompanying strong increase in chromatin-associated non-coding RNA levels and not related to a perturbing effect of actively transcribing polymerases.

Next, we sought to provide more insight into the molecular mechanisms by which strongly increased chromatin-associated RNA levels disrupt enhancer-promoter interactions. In this respect, it is of interest that RNA molecules have recently been implicated in the regulation of transcriptional condensates^58,59^. Current models predict that low levels of (short) RNA promote condensate formation, whereas high levels of (longer) RNA cause dissolution of condensates^58^. This suggests that increased chromatin-associated RNA levels in the context of INTS11 depletion could destabilize transcriptional condensates and thereby reduce enhancer-promoter interactions. Since Mediator is a key component of transcriptional condensates^16^, this hypothesis predicts that INTS11 depletion is associated with fewer or smaller Mediator clusters. To explore this possibility, we further engineered the HCT116 INTS11-dTAG cell line and labelled MED19 with a JF646-HaloTag (**Extended Data Fig. 4i**) to enable visualization of Mediator clusters using structured illumination microscopy (SIM). This enables live-cell super-resolution imaging, allowing us to observe clusters over a broad size range, including small clusters that cannot be resolved by conventional imaging. These experiments show that INTS11 depletion alone or in combination with CDK9 inhibition does not significantly alter the number of Mediator clusters per cell or their size distribution (**Fig. 4g,h, Extended Data Fig. 4j,k**). Therefore, based on these data, we cannot explain RNA-induced disruption of enhancer-promoter interactions by noticeable changes in Mediator clustering.

To better understand the functional consequences of RNA-mediated perturbation of enhancer-promoter interactions, we plotted changes in transcription as measured by TT-seq of the genes analyzed by MCC (**Fig. 4i**). When we stratify these in quartiles based on the extent to which enhancer-promoter interactions are decreased, we observe that the genes with the most perturbed enhancer interactions have significantly reduced transcription levels. Furthermore, the extent to which gene transcription is reduced scales with the extent to which interactions with enhancers are decreased. Together, these results show that gene loci at which enhancer-promoter interactions are most reduced upon INTS11 depletion show the largest decrease in mRNA transcription.

### CDK9 inhibition reduces chromatin-associated RNA levels and stabilizes enhancer-promoter interactions

Our targeted separation-of-function perturbations of the Integrator and Restrictor complexes have allowed us to demonstrate that strongly increased chromatin-associated non-coding RNA levels disrupt enhancer-promoter interactions. To better understand the general role of RNA in regulating enhancer-promoter interactions, we sought to test how reducing RNA levels on chromatin would affect enhancer-promoter interactions. To this end, we characterized the effect of 45 minutes of CDK9 inhibition by NVP2 without simultaneously depleting INTS11. To validate effective inhibition of CDK9 in these conditions, we performed high-resolution ChIP experiments for Pol II and calculated the Pol II pausing index^60^, which is increased following NVP2 treatment, indicating effective CDK9 inhibition (**Fig. 5a, Extended Data Fig. 5a-d**). As expected, chromatin-associated RNA-seq experiments show that this is associated with a decrease in chromatin-associated mRNA, uaRNA and eRNA levels (**Fig. 5b,c**). In addition, chromatin proteomics experiments show that CDK9 inhibition is associated with an increase in chromatin-bound NELF subunits and a decrease in chromatin-bound transcription elongation factors, whereas Integrator, Mediator, cohesin and CTCF are unchanged (**Fig. 5d**, **Extended Data Fig. 5e**).

**Fig. 5.**
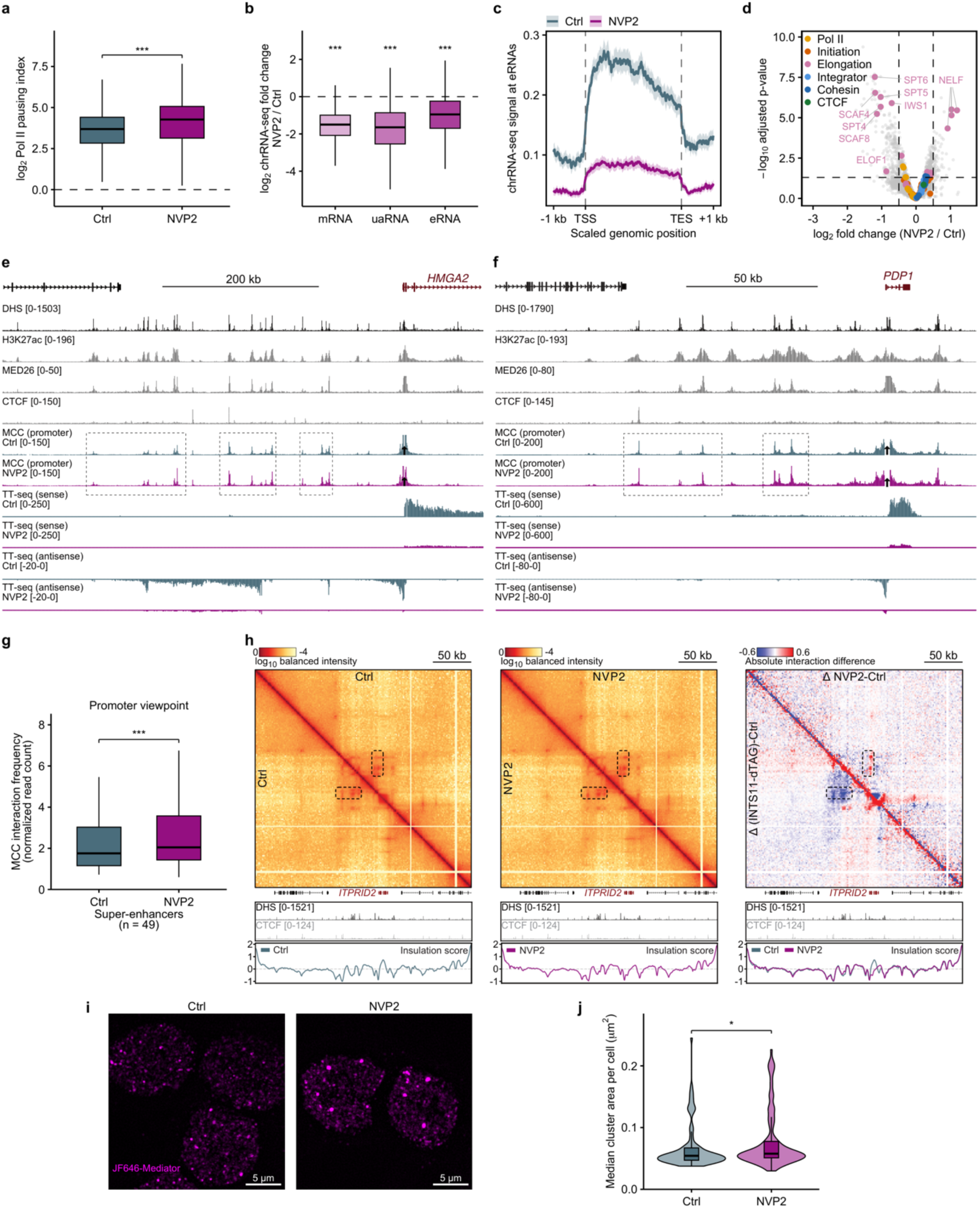
CDK9 inhibition reduces chromatin-associated RNA levels and stabilizes enhancer-promoter interactions. **a,** Pol II pausing indices (n = 6,136 genes) in Ctrl (grey) and upon NVP2 treatment for 45 min (magenta). Two-sided Wilcoxon signed-rank test. Box plots and tests as in Fig. 1. **b,** chrRNA-seq fold changes upon NVP2 treatment of mRNAs (n = 9,213), uaRNAs (n = 3,539) and eRNAs (n = 3,182). Box plots and tests as in Fig. 1. **c**, chrRNA-seq signal across eRNAs (n = 3,860), as in Fig. 1b. **d**, Chromatin proteomics upon NVP2 treatment (4,454 proteins); 76 proteins were enriched on, and 226 were lost from chromatin. Volcano plot as in Fig. 2j. **e**, MCC interaction profiles (mean of n = 3 biological replicates) from the HMGA2 promoter viewpoint and TT-seq in Ctrl (grey) and NVP2 (magenta). Tracks as in Fig. 1e. **f**, As in **e**, for the PDP1 locus. Tracks as in Fig. 1f. **g**, MCC interaction frequencies between promoter viewpoints and super-enhancers (n = 49) in Ctrl and NVP2. Box plots and tests as in Fig. 1g. P = 8.9 × 10^−7^. **h**, Tiled-MCC contact matrices of the ITPRID2 locus in Ctrl (mean of n = 3 biological replicates) and NVP2 (mean of n = 2 biological replicates) and differential matrices (top, ΔNVP2-Ctrl; bottom, (ΔINTS11-dTAG)-Ctrl), as in Fig. 1i. **i**, SIM of MED19 (Mediator) in INTS11-dTAG-MED19-Halo cells, as in Fig. 4g. **j**, Median MED19 cluster area per cell in Ctrl (n = 94 cells) and NVP2 (n = 103 cells), as in Fig. 4h. P = 0.038. *P < 0.05; ***P < 0.001.

To investigate whether reducing RNA levels on chromatin leads to changes in enhancer-promoter interactions, we performed additional MCC experiments. Interestingly, we observe that NVP2 treatment increases enhancer-promoter interactions (**Fig. 5e-g, Extended Data Fig. 5i,j**). Similar to the effect of increased RNA levels in the context of INTS11 depletion, we find that the effect of decreased RNA levels in the context of CDK9 inhibition is most pronounced on interactions between gene promoters and super-enhancers. To characterize changes in 3D genome folding induced by NVP2 treatment more comprehensively, we performed Tiled-MCC experiments. Direct comparison of the Tiled-MCC data in the context of CDK9 inhibition to INTS11 depletion further highlights the opposing effects of reducing and increasing RNA levels on 3D genome folding: CDK9 inhibition leads to increased enhancer-promoter interactions, whereas INTS11 depletion weakens these enhancer-promoter interactions (**Fig. 1i**, **Fig. 5h**). Taken together, we therefore conclude that CDK9 inhibition and the associated reduction in chromatin-associated RNA levels stabilize enhancer-promoter interactions.

To explore how CDK9 inhibition affects Mediator clustering, we performed additional live-cell SIM experiments. These experiments show that CDK9 inhibition leads to a significant decrease in the number of Mediator clusters per cell (**Fig. 5i, Extended Data Fig. 5k**), as well as a slight increase in the size of Mediator clusters (**Fig. 5i,j, Extended Data Fig. 5l**), which is consistent with previous live imaging experiments in the context of inhibition of transcription elongation^16,58^.

To further confirm that the increase in enhancer-promoter interactions upon CDK9 inhibition is related to a decrease in chromatin-associated RNA and not confounded by potential co-occurring transcription-mediated changes in loop extrusion^20,48–50^, we performed RAD21 ChIP experiments. These experiments show that cohesin occupancy is largely unchanged upon CDK9 inhibition (**Extended Data Fig. 5f-h**). Because ChIP experiments do not directly inform on the dynamic behavior of cohesin during its extrusion process and its potential interactions with Pol II, we used an orthogonal approach to test if and how cohesin interacts with Pol II and thereby further examine a potential confounding effect of CDK9 inhibition on cohesin-mediated loop extrusion. To this end, we performed biochemical reconstitution experiments with purified Pol II and cohesin complexes. Because NELF has also been implicated in enhancer-promoter interactions^21^ and because CDK9 inhibition leads to accumulation of paused Pol II and NELF on chromatin (**Fig. 5a,d**), we included this complex in our biochemical reconstitutions. Upon purifying all implicated protein complexes, we tested whether cohesin stably associates with any of the promoter-associated transcription complexes. Notably, we find that cohesin does not form a stable complex with either the core Mediator-bound pre-initiation complex or with paused Pol II elongation complexes (**Extended Data Fig. 6**). Together, these experiments therefore indicate that it is unlikely that changes in the distribution of these complexes on chromatin in the context of CDK9 inhibition (or INTS11 depletion) cause alterations in cohesin-mediated loop extrusion that underlie the observed changes in enhancer-promoter interactions.

## DISCUSSION

A key open question in the field of gene regulation concerns the molecular drivers of enhancer-promoter interactions. Research in this area has primarily focused on cohesin-mediated loop extrusion and protein-protein interactions involving transcription factors, co-activators and the transcription machinery. Here, we demonstrate that non-coding RNA molecules modulate long-range enhancer-promoter interactions, and thereby identify an additional regulatory principle of enhancer-promoter communication. More specifically, we discover that strongly increased chromatin-associated non-coding RNA levels upon INTS11 depletion disrupt enhancer-promoter interactions, whereas reduced RNA levels upon CDK9 inhibition are associated with stabilized enhancer-promoter interactions. The disruptive effect of RNA is predominantly visible at super-enhancers, which are more active and larger than typical enhancers, and therefore produce more and longer RNA molecules. Furthermore, a smaller increase in chromatin-associated non-coding RNA in the context of WDR82 depletion does not affect enhancer-promoter interactions. These two observations indicate a threshold response, whereby enhancer-promoter interactions are only disrupted when chromatin-associated RNA reaches a critical level.

What is the mechanism by which non-coding RNA modulates enhancer-promoter interactions? Changes in cohesin-mediated loop extrusion are unlikely to explain our observations, since cohesin occupancy is largely unchanged upon our perturbations and cohesin does not form stable complexes with the transcription machinery in biochemical reconstitution experiments. It remains in principle possible that the strong transcription units present at enhancers upon INTS11 depletion form an insulating boundary that interferes with interactions between additional enhancers and the gene promoter on opposite sides, a mechanism that has recently been proposed to underlie promoter competition^61^. This could in principle explain our observation that promoter interactions with distal enhancers are more strongly affected than proximal interactions, since there is usually a proximal enhancer cluster in between the more distal cluster and the gene promoter. However, in contrast to chromatin interaction data in the context of promoter competition^61^, our TMCC contact matrices do not show formation of insulating boundaries at upregulated enhancer clusters upon INTS11 depletion, indicating that increased insulation does not underlie the observed reduction in long-range enhancer-promoter interactions in this context.

Since many transcription factors and coactivators have been shown to directly bind or interact with RNA^62–65^, a second possible mechanism is that excessive non-coding RNA molecules could sequester these regulatory proteins and thereby reduce their occupancy on enhancers and promoters. Transcription factor binding at enhancers and promoters generally promotes accessibility and recruitment of Mediator^1,47^. Since INTS11 depletion does not reduce chromatin accessibility or Mediator recruitment at enhancers and promoters, and chromatin proteomics data show no general loss of sequence-specific transcription factors or Mediator from chromatin, our data do not support this mechanism, although we cannot fully rule out that INTS11 depletion may be associated with reduced occupancy of a subset of transcription factors or coactivators.

A third possible mechanism is that increased non-coding RNA transcription sequesters the transcription machinery towards non-coding elements, such as enhancers, at the expense of recruitment to gene promoters. However, WDR82 depletion increases non-coding transcription to a similar extent as combined INTS11 depletion and CDK9 inhibition, yet leaves enhancer-promoter interaction unchanged, indicating that these interactions are not caused by changes in the rate of non-coding RNA transcription. In addition, we do not observe reduced Mediator occupancy at gene promoters in INTS11 depletion conditions. Furthermore, even though Pol II is reduced at gene promoters upon INTS11 depletion, it is largely restored upon combined INTS11 depletion and CDK9 inhibition, and we continue to observe perturbed enhancer-promoter interactions in these conditions.

Enhancer-promoter interactions respond to changes in RNA concentration in a manner reminiscent of what has been reported for transcriptional condensates: strongly increased non-coding RNA levels exceeding a threshold disrupt transcriptional condensates, whereas reduced RNA levels stabilize these condensates^58,59^. A fourth possibility is therefore that RNA-dependent modulation of condensates underlies the observed changes in enhancer-promoter interactions. Mechanistically, this may be linked to the high negative charge density of the phosphate backbone of RNA molecules, which can modulate electrostatic interactions involved in condensate formation^58^. In this context, the modulation potential of eRNAs could be particularly strong, as they may be less heavily coated with proteins than mRNAs and thus expose more of their negatively charged backbone. Live imaging of Mediator shows an increase in the size of Mediator clusters upon CDK9 inhibition, which is in principle consistent with the increased enhancer-promoter interactions that we observe upon CDK9 inhibition. However, we failed to detect a reduction in cluster number or size upon INTS11 depletion (with and without CDK9 inhibition) that could explain the reduced enhancer-promoter interactions observed in these conditions. This indicates that the effect of RNA on enhancer-promoter interactions cannot be explained by global changes in Mediator clustering. However, because our experiments cannot resolve the subset of Mediator clusters at the individual loci at which we observe reduced enhancer-promoter interactions, it remains possible that more specific changes in a subset of Mediator clusters underlie the effect of RNA on enhancer-promoter interactions.

It is important to note that the disruptive potential of RNA is not necessarily restricted to biomolecular condensates. Therefore, a fifth possible mechanism is that at sufficiently high concentrations, RNA weakens discrete molecular interactions in solution by competing for protein-binding surfaces, by steric occlusion, or by changes in protein conformation and local electrostatic properties.

Taken together, our data indicate that excessive non-coding RNA does not generally affect the ability of enhancers and promoters to maintain accessibility or recruit proteins but disrupts the interactions between the molecules that are required for their 3D interactions in the nuclear space. The observed increased sensitivity of long-range enhancer-promoter pairs to RNA may reflect that they are more dependent on active mechanisms to facilitate interactions, because their greater genomic separation lowers their baseline contact probability^66^. The precise nature of these enhancer-promoter interactions and their potential relationship to transcriptional condensates remain to be determined in future studies using live-cell imaging of individual loci, which is beyond the scope of this study.

What is the function of RNA-mediated disruption of enhancer-promoter interactions? Since genes with the most reduced enhancer-promoter interactions show the greatest loss of transcription upon INTS11 depletion, it is likely that non-coding RNA accumulation provides a negative feedback mechanism, in which excessive RNA production at enhancers limits the interactions that drive transcription. Since increases in RNA are tolerated up to a certain threshold, this mechanism may be particularly important for super-enhancers, which produce large amounts of RNA and thus operate close to this threshold. RNA-mediated feedback regulation of enhancer-promoter interactions could therefore function as a mechanism to control the strong activation potential of super-enhancers, allowing cells to tune their long-range interactions without the need to reduce recruitment of regulatory proteins. At the same time, this implies that mechanisms that limit chromatin-associated RNA, such as Integrator-mediated termination and RNA degradation, are critical to allow for a strong gene activation potential by super-enhancers.

Together, our work demonstrates that non-coding RNAs are not merely a by-product of enhancer activity but act as functional regulatory molecules that control 3D enhancer-promoter interactions and transcription. Although previous observations have implicated (non-coding) transcription and/or RNA in 3D genome organization and gene regulation^27,51,63,67–70^, our work goes beyond previous studies, because our separation-of-function perturbations uncouple the functional contribution of the process of transcription that produces the RNA, the regulatory proteins that are recruited to *cis*-regulatory elements to initiate their transcription and the RNA molecules themselves. By establishing RNA as a regulator of enhancer-promoter interactions, our findings open new directions for investigating how its molecular properties and modifications modulate enhancer-mediated gene activation.

## METHODS

### Cell Culture

INTS11-dTAG human colorectal carcinoma HCT116 cells, in which both endogenous INTS11 alleles carry a C-terminal 2×HA-FKBP12^F36V (dTAG) tag^42^ (a gift from Steven West) were cultured in RPMI 1640 medium (Gibco, 21875034) supplemented with 10% FBS (Gibco, 10270106) and 1% GlutaMax (Gibco, 35050038) at 37 °C and 5% CO2. Cells were passaged every 3 days upon reaching ∼70-80% confluency by adding 0.05% trypsin (Gibco, 25300054) for 2 min.

For INTS11 depletion, dTAGv-1 (Tocris, 6914) was dissolved in DMSO. For CDK9 inhibition, NVP2 (MedChemExpress, HY-12214A) was dissolved in DMSO. Cells were seeded 36-40 h before treatment and treated at 80% confluency. Compounds were added directly to the medium: 0.5 μM dTAGv-1 for 90 min (INTS11-dTAG), 0.5 μM NVP2 for 45 min (NVP2), or 0.5 µM dTAGV-1 for 90 min with 0.5 µM NVP2 added for the final 45 min (INTS11-dTAG+NVP2). Control samples received an equivalent volume of DMSO for 90 min.

WDR82-mAID human colorectal carcinoma HCT116 cells^38^ (a gift from Steven West), in which both endogenous WDR82 alleles carry a C-terminal mini-AID (mAID) tag and OsTIR1 is expressed from a doxycycline-inducible cassette in the AAVS1 locus, were cultured under the same conditions. To induce OsTIR1, doxycycline (Sigma-Aldrich, D3447) was added to both control and treatment groups at the time of seeding (approximately 36 h before harvesting) to a final concentration of 1 µg/mL. WDR82 depletion was initiated by treating the cells with auxin (indole-3-acetic acid, IAA; Sigma-Aldrich, I2886; dissolved in DMSO) at a final concentration of 500 µM for 4 h. Control samples received an equivalent volume of DMSO.

Drosophila Schneider 2 (S2) cells, used as spike-ins, were obtained from DSMZ (ACC 130; RRID: CVCL_Z232) and cultured in Schneider’s Drosophila Medium (Gibco, 21720024) supplemented with 10% fetal bovine serum (Gibco, 10500-064) in a non-humidified incubator at 27°C, protected from light. For spike-in chromatin, 100 × 10⁶ S2 cells were crosslinked with 1% formaldehyde for 7.5 min and processed in the same way as the human chromatin.

All cell lines were verified to be free of mycoplasma contamination. Biological replicates are defined as independent cell cultures, grown separately and at different passage numbers, that were treated and processed as independent samples.

### TT-seq

TT-seq was performed in two biological replicates as previously described^43,71^ with modifications. For each sample, 80 × 10⁶ HCT116 INTS11-dTAG or WDR82-mAID cells were used. 4-thiouridine (4sU; Carbosynth, NT06186) was added at 0.5 mM for the final 5 min of treatment. Cells were lysed in TRIzol reagent (Invitrogen, 15596026), and 5 × 10⁶ Drosophila S2 cells labelled with 0.5 mM 4sU for 30 min were added to each sample. Six synthetic RNA spike-ins of 949-1,079 nt (three 4sU-labeled: Spike2, Spike4, Spike8; three unlabeled: Spike5, Spike9, Spike12)^43^ were added at 30 ng per 150 µg of total RNA. Total RNA was fragmented to below 10 kb on a Covaris S220 focused-ultrasonicator. Fragmented RNA was biotinylated with EZ-Link HPDP-Biotin (Thermo Fisher Scientific, 21341), and 4sU-labelled RNA was purified with the µMACS Streptavidin Kit (Miltenyi Biotec, 130-074-101) and cleaned up with the miRNeasy Micro Kit (Qiagen, 217084), including on-column DNase I digestion. Libraries were prepared from 100 ng of 4sU-labelled RNA with the Illumina Stranded Total RNA Prep, Ligation with Ribo-Zero Plus kit (9 PCR cycles). Fragment sizes were analyzed with a Fragment Analyzer (HS NGS Fragment Kit), and pooled libraries were sequenced on an Illumina NextSeq 2000 in paired-end mode (INTS11-dTAG, 2 × 65 bp; WDR82-mAID, 2 × 60 bp).

### MCC

MCC experiments were performed in three biological replicates per condition, following the improved MCC protocol^45^. After treatment, cells were harvested and fixed in batches of 50 million cells at a final concentration of 2% formaldehyde for 10 min. After quenching with glycine (0.15 M) and two washes with ice-cold PBS, cells were split into aliquots of 5 million and permeabilized with digitonin (0.005%, Sigma-Aldrich, D141). Each aliquot was immobilized on CUTANA™ Concanavalin A-conjugated paramagnetic beads (EpiCypher, 21-1401). Chromatin digestion was performed with MNase (NEB, M0247) in MNase buffer (50 mM Tris-HCl pH 7.5, 1 mM CaCl2) for 1 h at 37 °C. After confirming efficient digestion, DNA fragments were end-repaired in the presence of biotinylated dNTPs to generate ligation-ready, biotin-labeled fragments. Following ligation and de-crosslinking, DNA was extracted using the DNeasy Blood and Tissue Kit (Qiagen, 69504). Fragments were size-selected to remove mononucleosomal DNA using Mag-Bind TotalPure NGS beads (Omega Bio-tek, M1378). Samples were then sheared to sizes within a range of 180-200 bp using a Covaris S220 Focused-ultrasonicator (200 s; duty factor, 10%; peak incident power, 175 W; cycles per burst, 200). Prior to indexing, a streptavidin clean-up step was performed to remove unbound biotinylated dNTPs. Each sample was split into two reactions for indexing and subsequently pooled. Following size selection to remove primer dimers and a final streptavidin clean-up, 2ug of indexed DNA per replicate was used for the first hybridization reaction. Captured DNA was recovered using M-270 Streptavidin Dynabeads (Invitrogen, 63305) and amplified for 9 cycles. A second hybridization reaction was performed with the full volume of captured material, followed by a streptavidin pull-down and 7 cycles of PCR amplification. Final library quality was assessed with a Fragment Analyzer (HS NGS Fragment Kit), and sequenced on an Illumina NextSeq 2000 in paired-end mode (2 × 151 bp).

Capture oligonucleotides (120 nt) were designed using a python-based tool^72^ (https://oligo.readthedocs.io/en/latest/). The pool of oligonucleotides was ordered as a multiplexed panel of ssDNA 5ʹ-biotinylated oligonucleotides (IDT, xGen Custom Hybridization Capture Panels). Oligonucleotides were diluted to a final concentration of 2.9 nM for each hybridization reaction.

### Tiled-MCC

For Tiled-MCC, capture enrichment was performed using the Twist Standard Hyb and Wash Kit (Twist Bioscience, 116614), Twist Universal Blockers (Twist Bioscience, 100578), and Twist Binding and Purification Beads (Twist Bioscience, 100983), as described in the manufacturer’s protocol with slight modifications^45^. Equal amounts of uniquely indexed MCC libraries were pooled to yield 2 µg of total DNA per condition. Due to library availability constraints, three biological replicates were used for the control (DMSO) and two biological replicates for the dTAG, dTAG+NVP2 and NVP2 conditions. All libraries were pooled and split into hybridization tubes with a maximum of 2 µg of library pool per reaction. Samples were dried using a vacuum concentrator at 45°C and resuspended in 5 µL of Blocker Solution and 7 µL of Universal Blockers on ice. The Twist Hybridization Mix was pre-heated at 65°C, cooled to room temperature for 5 min, and combined with a previously designed Twist Custom Probe Panel^19^ and ddH2O to prepare the probe solution. The probe solution was incubated at 95°C for 2 min, transferred to ice or 4°C for 5 min, and left at room temperature for 5 min, while the resuspended library pool was simultaneously incubated at 95°C for 5 min and left at room temperature for 2 min. The probe solution was transferred to the library pool, 30 µL of Hybridization Enhancer was added, and the mixture was incubated for 16 h at 70°C on a thermocycler with the lid set to 85°C. Streptavidin Binding Beads were pre-washed three times with Binding Buffer, and the hybridization mixture was transferred directly to the beads and incubated for 30 min at room temperature at 600 rpm. The beads were washed once with Wash Buffer 1 and three times with Wash Buffer 2 at 48°C for 5 min. Following the final wash, beads were resuspended in 45 µL of ddH2O, split into two reactions, and subjected to post-capture PCR amplification using 2 × KAPA HiFi HotStart ReadyMix and Illumina Amplification Primers for 11 cycles. The PCR reactions were pooled, purified using 1.8 × Mag-Bind TotalPure NGS beads (Omega Bio-tek, M1378), and eluted in 100 µL of ddH2O. Final libraries were assessed with a Fragment Analyzer (HS NGS Fragment Kit), and sequenced on an Illumina NextSeq 2000 in paired-end mode (2 × 151 bp).

### ATAC-seq

Assay for Transposase-Accessible Chromatin using sequencing (ATAC-seq) was performed as described previously^19,73^. Three biological replicates were analyzed per condition. Briefly, 1.5 × 10^5^ cells were washed, split over two tubes and pelleted by centrifugation. Cell pellets were resuspended in ice-cold lysis buffer (10 mM Tris-HCl pH 7.5, 10 mM NaCl, 3 mM MgCl2, 0.1% Igepal CA-630) and incubated on ice for 3 min. Nuclei were washed once with cold PBS and resuspended in tagmentation mix (Illumina, 20034198). The tagmentation reactions were incubated for 30 min at 37 °C at 1,000 rpm. After DNA purification using the MinElute PCR purification kit (Qiagen, 28004), the tagmented samples were amplified using Nextera indexing primers and NEBNext High-Fidelity PCR Master Mix (NEB, M0541) for 11 cycles. Double size selection was performed with Mag-Bind TotalPure NGS beads (Omega Bio-tek, M1378) to remove primer dimers and fragments larger than 700 bp. Library quality was assessed with a Fragment Analyzer (HS NGS Fragment Kit), and sequenced on an Illumina NextSeq 2000 in paired-end mode (2 × 66 bp).

### Mediator and cohesin ChIPmentation

ChIPmentation for MED26 and RAD21 was performed in three biological replicates per condition as previously described^74,75^. Briefly, 1 M cells were fixed with 1% formaldehyde (28908, Thermo Scientific) for 10 min at room temperature, quenched with ice-cold glycine (Sigma-Aldrich, G7126) at a final concentration of 125 mM and washed two times with ice-cold PBS. Two-step lysis was performed by first isolating the nuclei with Farnham lysis buffer and then performing nuclear lysis with a 0.5% SDS lysis buffer. Chromatin was fragmented by a titrated amount of MNase (5.6 Kunitz) and then mildly sonicated (2% duty factor, 105 W peak incident power, 200 cycles per burst) for 1 min for MED26 and 12 min for RAD21. Following quality control of the fragmented chromatin profiles, 5 µg of sheared chromatin was added to a pre-incubated mixture of bead-bound antibodies (2 µg of target antibody; anti-MED26, Cell Signaling, 13641; anti-RAD21, Abcam, EPR22506-15; 1 µg of spike-in antibody; Active Motif, 61686), alongside with 50 ng spike-in chromatin, and incubated overnight at 4 °C with gentle rotation. After 16 h, samples were washed with stringent buffers and bead-bound DNA was tagmented using the Illumina Tagment DNA Enzyme and Buffer Kit. Following additional washes, samples were de-crosslinked overnight at 65 °C with proteinase K (0.2 mg/mL) and purified using Mag-Bind TotalPure NGS beads (Omega Bio-tek, M1378). The resulting DNA was amplified with the NEBNext High-Fidelity 2 × PCR Master Mix, assessed with a Fragment Analyzer (HS NGS Fragment Kit), and sequenced on an Illumina NextSeq 2000 in paired-end mode (2 × 66 bp).

### Pol II xChIP-seq

x-ChIP-seq for RPB1 was performed in three biological replicates per condition as previously described^76^, with modifications. For each sample, 40 × 10⁶ cells were used. After treatment, cells were crosslinked with 1% formaldehyde (16% methanol-free, Thermo Fisher Scientific, 28908) for 7.5 min at room temperature. Crosslinking was quenched with 125 mM glycine for 5 min. Cells were washed with PBS, scraped in ice-cold PBS, and pelleted at 300 × g for 5 min at 4°C; pellets were snap-frozen and stored at −70°C. All buffers, except the wash and elution buffers, were supplemented with a protease inhibitor cocktail (EDTA-free; Bimake, B14002) and a phosphatase inhibitor cocktail (APExBIO, K1015). Pellets were lysed in 5 mL Farnham Lysis Buffer (5 mM PIPES pH 8.0, 85 mM KCl, 0.5% (v/v) NP-40) for 10 min on ice, and nuclei were pelleted at 1,700 × g for 5 min at 4°C and washed with PBS. Nuclei were resuspended in 200 µL 1% SDS Lysis Buffer (50 mM Tris-HCl pH 8.0, 10 mM EDTA, 1% SDS) for 10 min at room temperature, diluted with 1.8 mL IP Buffer (20 mM Tris-HCl pH 8.0, 150 mM NaCl, 1% Triton X-100) supplemented with 5 mM CaCl2, and pre-warmed at 37°C for 2 min. Chromatin was digested with 25,000 U MNase (NEB, M0247S) at 37°C for 20 min, aiming for ∼80% mononucleosomes, and the reaction was stopped with EGTA and EDTA (10 mM and 5 mM final). Samples were sonicated in AFA milliTubes on a Covaris S220 focused-ultrasonicator (5% duty cycle, 140 W peak incident power, 200 cycles per burst, 720 s, 6°C, continuous degassing, water level 8) and clarified at 16,000 × g for 2 min at 4°C. Digestion was checked by TapeStation on an aliquot treated with RNase A (20 µg/mL, 30 min, 37°C) and Proteinase K (200 µg/mL, 2 h, 58°C) and purified (Qiagen PCR purification kit). For immunoprecipitation, 55 µg of HCT116 chromatin and 100 ng of Drosophila S2 spike-in chromatin were diluted in IP Buffer to a final 0.05% SDS in 1.4 mL. Chromatin was pre-cleared with Protein G Dynabeads (Invitrogen, 10004D; washed in PBS with 0.02% Tween) for 30 min at 4°C, and incubated overnight at 4°C with 10 µg anti-RPB1 antibody (Diagenode, C15200004), and cleared by centrifugation (16,000 × g, 2 min, 4°C). Immune complexes were captured on Dynabeads for 2 h at 4°C and washed for 5 min each at 4°C, once with TSE I (20 mM Tris-HCl pH 8.0, 150 mM NaCl, 2 mM EDTA, 1% Triton X-100, 0.1% SDS), four times with TSE II (as TSE I with 500 mM NaCl), once with Buffer III (10 mM Tris-HCl pH 8.0, 250 mM LiCl, 1 mM EDTA, 1% NP-40, 1% sodium deoxycholate), and three times with TE containing 50 mM NaCl. Chromatin was eluted in 50 µL elution buffer (0.1 M NaHCO3, 160 mM NaCl, 1% SDS), de-crosslinked overnight at 65°C, and treated with RNase A (20 µg/mL, 1.5 h, 37°C, after adjusting to 10 mM EDTA and 40 mM Tris-HCl pH 6.5) and Proteinase K (200 µg/mL, 2 h, 45°C). DNA was purified with the Monarch Oligonucleotide

Cleanup Kit (NEB, T1030) and quantified with a Qubit dsDNA HS Assay Kit. Libraries were prepared from 20 ng of DNA with the NEBNext Ultra II DNA Library Prep Kit (NEB, E7645) at half the recommended volumes, with an end-prep program adjusted for short fragments (30 min at 20°C, 60 min at 50°C, heated lid 60°C), and adapters diluted 18-fold. Adapter-ligated DNA was size-selected with AMPure XP beads (1.1 × and 0.6 ×), amplified with 10 PCR cycles and size-selected again (1.0 × and 0.4 ×). Fragment sizes were analyzed with a Fragment Analyzer (HS NGS Fragment Kit), and pooled libraries were sequenced on an Illumina NextSeq 2000 in paired-end mode (2 × 41 bp).

### Chromatin proteomics

Chromatin proteomics was performed in five biological replicates as previously described^76^, with modifications. For each sample, 12 × 10^6^ cells were used. Chromatin was isolated by cellular fractionation^77^. All buffers were supplemented with protease (cOmplete, Roche) and phosphatase (PhosSTOP, Roche) inhibitors. After treatment, cells were washed with ice-cold PBS, scraped from the plate, pelleted at 300 × g for 5 min at 4°C, and washed twice in DPBS. Cells were lysed in 400 µL Cell Lysis Buffer (10 mM Tris-HCl pH 7.4, 150 mM NaCl, 0.15% (v/v) NP-40) for 5 min on ice, overlaid on 1 mL Sucrose Buffer (10 mM Tris-HCl pH 7.4, 150 mM NaCl, 24% (w/v) sucrose), and centrifuged at 1,000 × g for 5 min at 4°C to pellet nuclei. Nuclei were washed in DPBS, resuspended in 250 µL Glycerol Buffer (20 mM Tris-HCl pH 7.4, 75 mM NaCl, 0.5 mM EDTA, 50% (v/v) glycerol, 1 mM DTT), and lysed by adding 250 µL Nuclear Lysis Buffer (10 mM Tris-HCl pH 7.4, 1 M urea, 300 mM NaCl, 0.2 mM EDTA, 7.5 mM MgCl2, 1% (v/v) NP-40, 1 mM DTT) for 2 min on ice. Chromatin was pelleted at 13,000 × g for 2 min at 4°C, and the nucleoplasmic supernatant was removed. The chromatin pellet was washed twice with Wash Buffer (10 mM Tris-HCl pH 7.4) and digested in 100 µL Chromatin Digest Buffer (50 mM Tris-HCl pH 7.4, 100 mM NaCl, 2.5 mM MgCl2, 2.5 mM CaCl2, 125 U Benzonase (Sigma-Aldrich, E1014), 800 U MNase (NEB, M0247S)) for 60 min at 4°C at 800 rpm. The digestion was stopped with 100 µL 2× SDS Buffer (100 mM Tris-HCl pH 7.4, 4% SDS), and samples were boiled at 99°C for 5 min. Insoluble material was pelleted at 10,000 × g for 10 min, and the solubilized chromatin extract was collected. 100 µg of protein from each chromatin extract was reduced with 10 mM DTT at 37 °C for 30 min and alkylated with 40 mM iodoacetamide at 37 °C for 30 min in the dark. Sera-Mag SpeedBeads (GE Healthcare, 45152101010250 and 65152105050250) were washed twice with water and added to the samples at a bead-to-protein ratio of 10:1 (w/w). Acetonitrile (ACN) was added to a final concentration of 50% (v/v), followed by incubation for 10 min at room temperature. After magnetic separation, the beads were washed three times with 90% ACN and once with 100% ACN, then resuspended in 50 mM triethylammonium bicarbonate (TEAB) containing 1 mM MgCl₂. The samples were incubated with 1 µL of Pierce Universal Nuclease (250 U; Thermo Fisher Scientific, cat. no. 88702) at 37°C for 3 h, followed by overnight digestion with sequencing-grade modified trypsin (1:20, enzyme-to-protein, w/w) at 37°C. The resulting peptides were collected, dried using a SpeedVac concentrator, and stored at −80°C until LC-MS/MS analysis (see below).

### chrRNA-seq

chrRNA-seq was performed in two biological replicates. For each sample, 12 × 10⁶ cells were used. After treatment, cells were scraped in ice-cold DPBS, 7.5 × 10⁶ Drosophila S2 cells were added per sample as a spike-in, and cells were pelleted at 300 × g for 5 min at 4°C. Chromatin was isolated by cellular fractionation^78^, with all buffers supplemented with a protease inhibitor cocktail (EDTA-free; Bimake, B14002). Cells were lysed in 400 µL Cell Lysis Buffer (10 mM Tris-HCl pH 7.4, 150 mM NaCl, 0.15% (v/v) NP-40) for 5 min on ice, overlaid on 1 mL Sucrose Buffer (10 mM Tris-HCl pH 7.4, 150 mM NaCl, 24% (w/v) sucrose), and centrifuged at 1,000 × g for 5 min at 4°C. Nuclei were washed in DPBS, resuspended in 125 µL Glycerol Buffer (NUN1; 20 mM Tris-HCl pH 8.0, 75 mM NaCl, 0.5 mM EDTA, 50% (v/v) glycerol, 1 mM DTT), and lysed by adding 1.2 mL Nuclear Lysis Buffer (NUN2; 20 mM HEPES pH 7.5, 1 M urea, 300 mM NaCl, 0.2 mM EDTA, 7.5 mM MgCl2, 1% (v/v) NP-40, 0.3% (v/v) Empigen, 1 mM DTT) with mixing for 10 min on ice. Chromatin was pelleted at 13,000 × g for 2 min at 4°C and washed with ice-cold DPBS. The chromatin pellet was resuspended in 95 µL Chromatin Digest Buffer (1× TURBO DNase Buffer, 0.1 U/µL TURBO DNase (Thermo Fisher Scientific, AM2238), 1 U/µL murine RNase inhibitor (NEB, M0314)) and incubated at 37°C at 1,400 rpm for 10 min to release chromatin-associated RNA; 5 µL Proteinase K (NEB, P8107) was added for a further 5 min at 37°C. RNA was extracted with TRI Reagent (Zymo, R2050) and phenol-chloroform and purified with the RNA Clean & Concentrator-25 kit (Zymo, R1017), and 20 ng of the six synthetic RNA spike-ins (see TT-seq section) were added per sample. After DNase I treatment (NEB, M0303) and a second clean-up, RNA integrity was checked on a TapeStation. Libraries were prepared from 250 ng of RNA with the Illumina Stranded Total RNA Prep, Ligation with Ribo-Zero Plus kit at half the recommended volumes, with a 5 min fragmentation at 94°C, an additional bead size selection and 7 PCR cycles. Fragment sizes were analyzed with a Fragment Analyzer (HS NGS Fragment Kit), and pooled libraries were sequenced on an Illumina NextSeq 2000 in paired-end mode (2 × 60 bp).

### Protein purification

General transcription factors (TFIIA, TFIIB, TFIIE, TFIIF, TBP, and TFIIH), Sus scrofa-derived RNAPII and core Mediator were expressed and purified as previously described^79–82^. For the paused elongation complex, DSIF and NELF were expressed and purified as previously described^83,84^. Factors used for assembly of the active elongation complex were expressed and purified as previously described^85^. Expression and purification of the cohesin complex containing SMC1A, SMC3, RAD21, STAG1 and NIPBL was performed as previously described^86^, with modifications. Briefly, the complex was co-expressed as three plasmids using the Bac-to-BacTM Baculovirus Expression System (Thermo Fisher). The plasmids encoded for C-terminally His6-MBP-tagged RAD21 (R172A, D279A, R450A mutations), untagged SMC3 and SMC1A, and NIPBL (C-terminal His6-MBP-tag) together with STAG1 (N-terminal His6-MBP tag). 2 liters of Hi5 cells (Thermo Fisher) expressing all three plasmids were centrifuged, resuspended and snap frozen in buffer A1 (300 mM NaCl, 20 mM Na-HEPES pH 7.5 (4°C), 10 % glycerol, 0.02 % Tween-20, 1 mM DTT, 0.284 μg/mL leupeptin, 1.37 μg/mL pepstatin A, 0.17 mg/mL PMSF and 0.33 mg/mL benzamidine). Cells were thawed, lysed by sonication, and the lysate cleared by centrifugation at 70,000 × g for 45 min, filtered, and applied to an XK 16/20 column packed with amylose resin. The column was washed with 4 column volumes (CV) of buffer A1 and 2 CV of buffer A2 (75 mM NaCl, 20 mM Na-HEPES pH 7.5 (4°C), 10 % glycerol, 1 mM DTT, 0.284 μg/mL leupeptin, 1.37 μg/mL pepstatin A, 0.17 mg/mL PMSF and 0.33 mg/mL benzamidine) before elution directly on a 5 mL HiTrap-Heparin HP column (Cytiva) with buffer A2 supplemented with 50 mM maltose. After washing the Heparin column with 8 CV of buffer A2, the cohesin complex was eluted with a gradient over 10 CV into buffer B2 (2 M NaCl, 20 mM Na-HEPES pH 7.5 (4°C), 10 % glycerol, 1 mM DTT). Peak fractions were analyzed by SDS-PAGE and pooled for subsequent TEV cleavage overnight. The digested complex was then concentrated using a pre-equilibrated 100 kDa MWCO Amicon Ultra Centrifugal Filter (Milipore) and applied to a Superose6 10/300 size exclusion column equilibrated in 200 mM NaCl, 20 mM Na-HEPES pH 7.5 (4°C), 10 % glycerol, and 1 mM TCEP. Peak fractions were analyzed by SDS-PAGE and those containing the cohesin complex were pooled, concentrated in pre-equilibrated 100 kDa MWCO Amicon Ultra Centrifugal Filters (Milipore) and snap frozen in liquid nitrogen for storage in-70°C.

### Complex assembly and analytical gradient ultracentrifugation

The pre-initiation complex with core Mediator was assembled on a shortened AdMLP promoter sequence (template:5’- CAGACAGCGATGCGGAAGAGAGTGAGGACGAACGCGCCCCCACCCCCTTTTATAGCCCCCCTTC-3’; non-template: 5’- GAAGGGGGGCTATAAAAGGGGGTGGGGGCGCGTTCGTCCTCACTCTCTTCCGCATCGCTGTCTG-3’) similar as previously described^81^. 2 µM TBP, TFIIA and TFIIB were assembled on the 0.5 µM annealed DNA scaffold. In parallel, 0.5 µM RNAPII and 2 µM TFIIF were pre-mixed, as well as 1 µM TFIIH-core with 1 µM TFIIH CAK. Full assembly was achieved by adding the pre-formed RNAPII-TFIIF complex, 1 µM TFIIE, and the pre-assembled TFIIH, with 2-minute incubation steps at 25°C between the individual additions. Lastly, 0.6 µM human core Mediator was added to form the PIC-cMed complex and the reaction was brought to a final volume of 30 µL in 100 mM KCl, 20 mM Na-HEPES pH 7.5 (at 4°C), 2 mM MgCl2, 5 % glycerol, and 2 mM TCEP. Depending on the tested complex, 2 µM cohesin was added together with the Mediator complex. For final assembly, the sample was incubated at 25°C at 400 rpm. Afterwards, aggregations were removed by centrifugation (21,000 × g, 5 min) and the reconstituted complex was applied to a 10-50% glycerol gradient formed in 2 mL centrifugation tubes (Beckmann Coulter) using a BioComp Gradient Master 108 (BioComp Instruments) by mixing a 10 % and 50 % glycerol solution (10/50 % glycerol (v/v), 100 mM KCl, 20 mM Na-HEPES pH 7.5 (at 4°C), 2 mM MgCl2, and 1 mM TCEP). The gradient was then centrifuged for 3 h at 259,000 × g and 4°C (Optima MAX-XP, Beckman Coulter), harvested in 100 µL fractions, and analyzed by SDS-PAGE.

Assembly of the paused elongation complex (PEC) with and without cohesin was performed on a DNA-mismatch bubble (template: 5’- TCAAGTACTTAAGCCTGGTCTATACTAGTA-3’, non-template: 5’-TACTAGTATTCTAGTATTGAAAGTACTTGA-3’) with an RNA primer (5’-rCrArGrArCrCrArGrGrC-3’). Assembly of the PEC was achieved by incubating 0.75 µM of RNA-template-DNA duplex with 0.5 µM of RNAPII for 10 min at 30°C, before adding 1.25 µM of the non-template DNA and 2 µM DSIF and NELF in 10 min of incubation intervals. 2 µM cohesin was optionally added as a last component before adjusting the reaction to a final volume of 30 µL (100 mM NaCl, 20 mM Na-HEPES pH 7.5 (at 4°C), 2 mM MgCl2, 4 % glycerol and 1 mM DTT) and incubation for 30 min at 30°C. The assembled complex was then applied to a 10-30 % sucrose gradient formed in 2 mL centrifugation tubes (Beckman Coulter) using a BioComp Gradient Master 108 (BioComp Instruments) by mixing a 10 % and 30 % sucrose solution (10/30 % sucrose, 100 mM NaCl, 20 mM Na-HEPES pH 7.5 (at 4°C), 2 mM MgCl2, 4 % glycerol (v/v), and 1 mM DTT). The gradient was centrifuged for 3 h at 259,000 × g and 4°C (Optima MAX-XP, Beckman Coulter), harvested in 100 µL fractions, and analyzed by SDS-PAGE.

The active elongation complex was assembled in a final volume of 35 µL containing 115 mM NaCl, 20 mM Na-HEPES pH 7.5 (at 4°C), 2 mM MgCl2, 4.4 % glycerol (v/v), and 1 mM DTT as previously described^85^. In brief, DNA oligos were annealed (template: 5’-ATCACTGTCGCGGCCCTTGTGTTCAGGAGCCAGCAGGGAGCTGGGAGCAAAAAGACTTAGAAAAATT

TTT-3’; non-template: 5’- AAAAATTTTTCTAAGTCTTTTTGCTCCCAGCTCCCTGCTGGCTCCGAGTGGGTTCTGCCGCGACAGTGA

T-3’). 0.75 µM of it was mixed with 1.35 µM RNA primer (5’- rUrUrArArGrGrArArUrUrArArGrUrCrGrUrGrCrGrUrCrUrArArUrArArCrCrGrGrArGrArGrGrGrArA rCrCrCrArCrU-3’) and incubated on ice for 10 min. 0.9 µM of RNAPII was added. After 10 min of incubation on ice, 1.8 µM of DSIF, SPT6, IWS1, RTF1, PAF1c, ELOF1 and 1 µM P-TEFb were added together with 1 µM ATP and buffer for a total of 35 µL, followed by incubation at 30°C for 15 min. The assembled complex was then applied to a 10-30 % sucrose gradient formed in 2 mL centrifugation tubes (Beckmann Coulter) using a BioComp Gradient Master 108 (BioComp Instruments) by mixing a 10 % and 30 % sucrose solution (10/30 % sucrose, 100 mM NaCl, 20 mM Na-HEPES pH 7.5 (at 4°C), 2 mM MgCl2, 4 % glycerol (v/v), and 1 mM DTT). The gradient was then centrifuged for 3 h at 259,000 × g and 4°C (Optima MAX-XP, Beckman Coulter), harvested in 100 µL fractions, and analyzed by SDS-PAGE.

### Western blotting

Following treatment, cells were harvested and washed once with ice-cold PBS. The cell pellet was resuspended in 50 µL of RIPA buffer (50 mM Tris-HCl (pH 8.0), 150 mM NaCl, 1% NP-40, 0.5% sodium deoxycholate, 0.1% SDS, 2 mM MgCl₂), supplemented with 1 × protease and phosphatase inhibitors and 2 µL/mL Benzonase (Sigma, E1014). The cell suspension was incubated with agitation for 1h at 4°C and then centrifuged at maximum speed at 4°C for 15 min. The concentration of the protein extract was measured using the Pierce BCA Protein Assay Kit (Thermo Fisher Scientific). 10-20 µg of protein extract were diluted in 6X Laemmli SDS sample buffer (Thermo Fisher Scientific) with 50 mM DTT (Carl Roth, 6908.3) and denatured for 5 min at 95°C. Proteins were separated on a NuPAGE 4-12% Bis-Tris gel (Invitrogen, NP0321) at 200 V for 30-40 min and blotted to a PVDF membrane. Each membrane was stained with Ponceau S to assess protein loading and transfer. After destaining, membranes were blocked for 1 h at room temperature using 5% nonfat milk in PBS containing 0.05% Tween-20 (PBS-T). The membrane was then briefly washed with PBS-T and incubated overnight at 4°C with the primary antibody. HA-tagged INTS11 was detected using rat monoclonal anti-HA (clone 3F10; Sigma-Aldrich, 11867431001; 1:2,000), and mAID-tagged WDR82 was detected using rabbit polyclonal anti-mAID (Proteintech, 28209-1-AP; 1:1,000). Halo-tagged MED19 was detected using mouse monoclonal anti-Halo (Promega, G9211; 1:1,000). Rabbit monoclonal anti-Cyclophilin B (clone EPR12703(B); Abcam, ab178397; 1:5,000) was used as a loading control. Rabbit polyclonal anti-Histone H3 (Abcam, ab21054; 1:4,000) was used as loading control too. Following primary-antibody incubation, membranes were washed three times for 5 min in PBS-T containing 1% nonfat milk, then incubated for 1 hour at room temperature with HRP-fused secondary antibody diluted in 1% nonfat milk in PBS-T (1:4,000; goat anti-rabbit (Abcam, ab205718), goat anti-mouse (Abcam, ab205719), or goat anti-rat (Sigma-Aldrich, A9037)), followed by three additional 10-min washes. Signal was detected using the Pierce™ ECL Western Blotting-Substrate kit (Thermo Fisher Scientific) and acquired with a ChemiDoc Imaging System (Bio Rad).

### Plasmid Construction

The MED19-Halo donor template for homology-directed recombination (HDR) was constructed by modifying the pUC19 cloning vector (New England Biolabs, N3041S). To achieve an N-terminal fusion, 250 bp homology arms (IDT) were designed to flank the start codon of the endogenous MED19 locus (hg38). The Halo construct (a gift from Ibrahim Cissé), followed by a short flexible (Gly₄Ser)₃ linker, and the homology arms were integrated into the pUC19 vector via a four-fragment Gibson assembly reaction (New England Biolabs, E2611S). Prior to that, the pUC19 backbone was linearized using EcoRI-HF (New England Biolabs, R3101S) and dephosphorylated. Primers were designed to achieve 25-30 bp overlap among the fragments. Assembly was performed with a 2:1 insert-to-vector molar ratio using 0.05 pmol of each DNA fragment. Colonies were picked and screened via colony PCR and validated by Sanger sequencing. High-quality supercoiled plasmid was isolated using the QIAGEN Plasmid Maxi Kit (12163) for subsequent nucleofection.

### Cell line generation

Endogenous knock-in was performed using the Alt-R HDR-mediated CRISPR-Cas9 system (IDT). CRISPR RNA (crRNA) targeting the N-terminus of MED19 was designed using the Alt-R CRISPR-Cas9 HDR Design Tool, prioritizing high on-target and low off-target scores and minimal distance between the cleavage and knock-in sites. The crRNA was designed with the spacer sequence 5ʹ-GGGTACGATGGAGAATTTCA-3ʹ. The genomic target was selected immediately upstream of the CGG protospacer adjacent motif (PAM). Ribonucleoprotein (RNP) complexes were assembled by annealing crRNA and tracrRNA (1.2:1 molar ratio, 50 μM final concentration) at 95°C for 5 min, followed by complexing with Alt-R S.p. HiFi Cas9 Nuclease (62 μM; IDT) for 10 min at room temperature. The HiFi Cas9 variant was utilized to minimize off-target effects.

Nucleofection was conducted using the 4D-Nucleofector System (Lonza) with the SE Cell Line 4D-Nucleofector kit [EN-113]. For each reaction, 2×10^5^ cells were resuspended in SE solution and combined with the RNP complex and the supercoiled MED19-Halo donor template. After nucleofection, cells were cultured in medium supplemented with 0.5 μM Alt-R HDR Enhancer V2 (IDT, 10007910). Monoclonal cell lines were established 48 h post-nucleofection via fluorescence-activated cell sorting (FACS) to isolate single Halo-positive cells. For FACS preparation, MED19 was labeled with 10 nM JF-646 Halo ligand (Promega, GA112A) for 30 min at 37 °C in the cell culture incubator. The cells were then washed three times with warmed PBS and incubated for 20 min in growth media. Labelled cells were centrifuged and resuspended in FACS buffer (1% FBS in 1x PBS buffer), then strained through a 35 μm sieve (Falcon, 2235). Single cell sorting was performed using the FACS Aria III using 633nm laser excitation. Genotyping PCR allowed the selection of homozygous MED19-Halo cells, which were further validated with Sanger Sequencing and Western Blot.

### Structured Illumination microscopy (SIM)

Two days prior to imaging, 2×10^5^ HCT116-INTS11-dTAG-MED19-Halo cells were seeded onto poly-L-ornithine (PLO)-coated glass-bottom dishes (MatTek Corporation, P35G-1.5-20-C). On the morning of the imaging session, cells were incubated with the JF-646 HaloTag ligand as described above. Experimental treatments were applied following MED19 labeling. Each condition was imaged sequentially following its respective treatment duration, with fields of view acquired from non-adjacent regions across the dish to minimize sampling bias.

Live-cell SIM was performed using a Zeiss Elyra 7 lattice SIM system equipped with a Plan-Apochromat 63×/1.46 Oil objective and a PCO.edge 4.2 CLHS sCMOS camera. Samples were illuminated with a 642-nm laser at 50% power (corresponding to an irradiance of 500 W cm^-2^ in widefield mode) with an exposure time of 100 ms. Z-stacks spanning a depth of 1.5 μm were acquired using a 32 μm (642 nm channel) grating pattern with 13 phase steps. Raw images were reconstructed using the standard Zeiss SIM processing algorithm with the ‘Scale to Raw Image’ option enabled to normalize intensity variations between fields of view while preserving relative intra-field intensities. Maximum intensity projections (MIPs) were generated and processed in batch mode using the pixel classification workflow in ilastik^87^. The resulting segmentation maps were imported into Fiji/ImageJ^88^, where cluster counts and surface areas (μm²) were quantified using the ‘Analyze Particles’ plugin. Clusters with areas of 30-500 pixels were retained, using a pixel size of 31 × 31 nm (0.029-0.481 μm²). Median cluster area was calculated per cell. Cells without retained clusters contributed zero to cluster counts and were excluded from median-area analyses. Data visualization and statistical plotting were performed using R. Per-cell values were compared with Ctrl using two-sided Wilcoxon rank-sum tests, with Benjamini-Hochberg correction across treatment comparisons within each panel.

### Analysis of high-throughput sequencing data

All sequencing data were demultiplexed with BCL Convert (Illumina; v4.4.6) and subsequently quality-checked with FastQC (Babraham Institute).

### TT-seq and chrRNA-seq data preprocessing and normalization

For all samples, adapters (ACTGTCTCTTATACACATCT) and bases below Q20 were removed with Cutadapt (v5.0), and read pairs with either mate shorter than 16 bp were discarded; for the INTS11-dTAG TT-seq libraries the first base of each mate was additionally clipped. Paired-end reads were mapped with STAR (v2.7.11b) first to the human rDNA repeat (GenBank U13369.1); unmapped pairs were then mapped to the GRCh38 (hg38) genome merged with the Drosophila melanogaster (dm6) genome and the synthetic RNA spike-in sequences, with a maximum 2% mismatches [mapping parameters: --outFilterMismatchNoverLmax 0.02 --outFilterMultimapNmax 1 --outFilterMatchNmin 16 --outFilterScoreMinOverLread 0 -- outFilterMatchNminOverLread 0 --alignIntronMax 500000]. Duplicate pairs were marked with Picard MarkDuplicates (v3.3.0) and excluded from analysis. SAMtools (v1.21) was used to quality filter alignments, whereby alignments with MAPQ smaller than 10 (-q 10) were skipped and only proper pairs (-f 0x2) were selected. Strand-specific coverage was taken from the aligned blocks of both mates, with the second mate defining the strand.

A spike-in normalization strategy was used to obtain a per-library size factor. A Drosophila size factor was estimated from read counts on Drosophila genes, and a synthetic-RNA size factor from read counts on the spike-in sequences, each as a DESeq2^89^ median-of-ratios factor. For TT-seq, the synthetic-RNA factor used only the three 4sU-labelled spike-ins; for chrRNA-seq, all six spike-ins were used. The size factor applied to each library was the geometric mean of its Drosophila and synthetic-RNA factors, rescaled to a geometric mean of 1 across the libraries of a dataset. Coverage and read counts divided by this factor are referred to as spike-in-normalized, and merged condition tracks are the arithmetic mean of the spike-in-normalized replicate coverages.

### MCC analysis

MCC data were analyzed using the C-MAP pipeline^45^. Briefly, raw reads underwent PCR deduplication via BBMap (version 38.18), adapter trimming (AGATCGGAAGAGCACACGTCTGAACTCCAGTCA and AGATCGGAAGAGCGTCGTGTAGGGAAAGAGTGT) with Trim Galore (version 0.6.10), and read merging via FLASH (version 1.2.11) prior to BWA-MEM alignment (version 0.7.12). Alignments were partitioned by viewpoint and filtered to retain reads containing a capture fragment and associated interacting reporter fragment. PCR duplicates—defined as reads with identical or near-identical alignments, allowing a 2 bp wobble at segment ends—were removed. Final interaction profiles were normalized for an equal number of interactions in cis and plotted as BigWig tracks using bedtools (version 2.31.1) and represent the mean of the biological replicates. Peaks were called and annotated as described previously^75^. Briefly, MCC peaks were annotated as promoters or enhancers based on their overlap with relevant genomic features. Promoters were defined by overlap with an ATAC-seq peak located within ±500 bp of the TSS of an expressed gene. In the remaining peaks, enhancers were defined by overlap with ATAC-seq and H3K27ac ChIP-seq signal and absence of CTCF ChIP-seq signal. Super-enhancers were defined as clusters of enhancers with H3K27ac and Med26 ChIP-seq enrichment. Normalized interaction frequencies with annotated region were quantified by calculating the mean signal of the top 70% of BigWig coverage values using a custom Python script. Enhancers more than 100 kb from their promoter were selected based on the distance between the center of the promoter and the enhancer. Treatments were compared with control by paired two-sided Wilcoxon signed-rank tests (exact for n ≤ 50), and P values were adjusted with the Benjamini-Hochberg method over the MCC comparisons shown in the same figure (for example, the five comparisons in Fig. 1g). For the comparison of proximal and distal enhancer interactions within individual loci (Figs 1h, 3h), at each promoter with at least two interactions with either super-enhancers or typical enhancers, the log2(treatment/control) change of the most distal interaction minus that of the most proximal was calculated and tested against zero. Data visualization and statistical plotting were performed using R.

### Tiled-MCC analysis

Tiled-MCC data were analyzed using the C-MAP pipeline^45^ as described above for MCC, using-tiled mode. To construct contact matrices for Tiled-MCC, reads containing at least one ligation junction within the tiled regions were retained. For each tiled region, ligation junctions were binned to generate raw contact matrices via cooler (v0.10.2), followed by matrix balancing using cooler balance. Multi-track genomic visualizations and insulation analyses were performed in Python (v3.9.18) using the cooler (v0.10.2), matplotlib (v3.8.2), and cooltools (v0.7.1) packages. Insulation scores were computed using cooltools.insulation with a sliding window size of ten times the bin resolution (ignoring the first two diagonals). Annotation tracks (DHS and CTCF ChIP-seq) were extracted using pyBigWig and aligned with the tiled region.

### ChIPmentation and ATAC-seq analysis

Raw paired-end reads were processed by trimming adapters (TCGTCGGCAGCGTCAGATGTGTATAAGAGACAG and

GTCTCGTGGGCTCGGAGATGTGTATAAGAGACAG) and low-quality bases (Phred score < 25) using Trim Galore (v0.6.10), followed by alignment to the human (hg38) and Drosophila (dm6) reference genomes using Bowtie2 (v2.4.4). After sorting and indexing with SAMtools (v1.13), duplicates were marked via Picard MarkDuplicates (v2.26) and removed alongside ENCODE-defined blacklist regions using bedtools (v2.30). Genome-wide coverage tracks were generated using the bamCoverage function from deepTools^90^ (v3.5.1) with a 10 bp bin size and 30 bp smoothing. MED26 tracks were normalized with a spike-in scale factor derived from the number of properly paired, deduplicated reads mapped to the D. melanogaster (dm6) genome. ATAC-seq and RAD21 tracks were normalized for sequencing depth (reads per kilobase per million mapped reads, RPKM). To generate representative tracks for each condition, BigWig files of individual replicates were merged by calculating the mean signal per bin using the bigwigAverage function.

Peaks were called with MACS3^91^ (v3.0.3) in paired-end mode (-f BAMPE-g hs-q 0.01--keep-dup all--call-summits). For each condition, the reads of all replicates were combined, and peaks were called once. The peaks of all conditions were then merged into one peak set per assay (union of all conditions), and peaks on chrM and non-canonical contigs were removed. The RAD21 peak set was further filtered for peaks overlapping RAD21 peaks in ENCODE data for HCT116 cells (ENCFF391AAM). Each peak was anchored at the median summit across the conditions in which it was called. Peaks were classed as promoter (overlapping TSS −250 to +50 bp of the major isoform of a protein-coding gene), super-enhancer (SE) or typical enhancer (TE) (defined as described in MCC analysis), or other. For differential analysis, reads were counted in 501-bp windows centered on the peak anchors (summarizeOverlaps; paired-end, unstranded). For ATAC-seq, the two technical libraries of each biological replicate were summed. Each assay was fitted in DESeq2 (design ∼ condition, DMSO as reference), with size factors from the in-window read totals (ATAC-seq, RAD21) or from dm6-aligned reads (MED26), each divided by its geometric mean. Log2 fold changes were not shrunk, independent filtering was disabled, and significant changes were defined as peaks with Benjamini-Hochberg-adjusted P < 0.05. One RAD21 NVP2 library (replicate 3) was excluded from the analysis as an outlier, based on a deviating low fraction of reads in peaks (5.7%) and a size factor of 0.59. In MA plots, SE features were peaks in MCC-targeted SE regions: SE-class peaks plus non-promoter peaks overlapping the body of an SE eRNA. Profiles and heatmaps were computed as described in Metagene profiles and heatmaps.

### RNA Pol II xChIP-seq data preprocessing and normalization

For all samples, adapters (AGATCGGAAGAGCACACGTCTGAACTCCAGTCA and AGATCGGAAGAGCGTCGTGTAGGGAAAGAGTGT) and bases below Q20 were removed with Cutadapt (v5.0;--nextseq-trim 20), and read pairs with either mate shorter than 16 bp were discarded. Reads were mapped with Bowtie 2 (v2.5.4;--very-sensitive --no-unal --no-mixed --no-discordant) to the GRCh38 (hg38) genome merged with the Drosophila melanogaster (dm6) genome. SAMtools (v1.21) retained primary, uniquely mapped proper pairs with MAPQ ≥ 10 (-q 10 -F 0x4 -F 0x100 -F 0x800). Duplicates were marked with Picard MarkDuplicates (v3.3.0) and retained, and reads overlapping the ENCODE blacklist (ENCFF356LFX) were removed with BEDTools (v2.31.1). For quantification, only fragments shorter than 70 bp were kept, and signal was taken as their per-base coverage. Coverage was normalized to a Drosophila spike-in size factor, calculated as the number of proper pairs mapping to the Drosophila chromosomes divided by the smallest such count across libraries. Merged condition tracks represent the arithmetic mean of the spike-in-normalized replicate coverages.

### Proteomics data analysis and visualization

LC-MS/MS analysis was performed on an Orbitrap Exploris 480 mass spectrometer (Thermo Fisher Scientific) coupled to a Vanquish Neo UHPLC system (Thermo Fisher Scientific). The UHPLC column setup consisted of a PepMap Neo C18 trap cartridge (5 µm particle size, 300 µm × 5 mm; Thermo Fisher Scientific) and a custom in-house-packed 34 cm C18 analytical column (75 µm inner diameter; ReproSil-Pur 120 C18-AQ, 3 µm particle size; Dr. Maisch GmbH). Peptides were separated at a flow rate of 300 nL/min using a gradient of 1-5% solvent B (0.08% (v/v) formic acid and 80% (v/v) acetonitrile) in 0.1 min, 5-10% B in 3 min, 10-30% B in 76.9 min, 30-44% B in 13 min, 44-55% B in 6 min, and 55-99% B in 1 min, followed by 99% B for 10 min. Solvent A consisted of 0.1% (v/v) formic acid in water. The total run time was 110 min. Mass spectrometry data were acquired in data-independent acquisition (DIA) mode using 40 variable isolation windows covering an m/z range of 350-1650. The Orbitrap resolution was set to 120,000 for MS1 and 30,000 for MS2. The normalized automatic gain control (AGC) targets were set to 300% for MS1 and 1,000% for MS2, with maximum injection times of 20 ms and 55 ms, respectively. Fragmentation was performed using higher-energy collisional dissociation (HCD) with a normalized collision energy (NCE) of 30%. Each sample was injected twice as a technical replicate. Peptide preparation and mass-spectrometry acquisition were carried out at the Bioanalytical Mass Spectrometry facility, Max Planck Institute for Multidisciplinary Sciences. Raw mass-spectrometry files were analyzed using DIA-NN (version 2.2.0, Academia) against the UniProt human reference proteome (UP000005640, June 2026; 20,652 entries), supplemented with common contaminants. An in silico predicted spectral library was generated. Trypsin digestion with up to one missed cleavage was specified. Carbamidomethylation of cysteine was set as a fixed modification and methionine oxidation as a variable modification. Peptide and protein identifications were filtered at a false discovery rate (FDR) of 1%. The protein-group quantity matrix was used for downstream analysis. The protein intensities of the two injections were averaged for each sample. Protein groups with more than 20% missing values across samples were removed, retaining 4,454 of 5,028 protein groups. The remaining intensities were log2-transformed and median-normalized across samples; missing values were not imputed. Differential abundance was assessed with limma. A linear model was fitted with condition as the primary factor and batch as a covariate; empirical Bayes moderation was applied, and P values were corrected for multiple testing with the Benjamini-Hochberg method. Three contrasts were tested: dTAG, NVP2, and dTAG+NVP2, each against DMSO. Proteins with an adjusted P < 0.05 and a |log2 fold change| > 0.5 were considered to be significantly changed. No batch correction was applied to the data shown. For plotting, protein groups were assigned to complexes by gene name: Pol II (POLR2A-L); Integrator (INTS1-15); initiation (the general transcription factors TFIIA, TFIIB, TFIID (TAF1-8 and TAF13), TFIIE, TFIIF and TFIIH, and all Mediator subunits including the CDK8 kinase module); elongation (NELF, DSIF, P-TEFb (CDK9, CCNT1), FACT, SPT6, the PAF1 complex, ELOF1, TFIIS, IWS1, SCAF4 and SCAF8); cohesin (SMC1A, SMC3, RAD21, STAG1, STAG2, NIPBL, MAU2, WAPL); and CTCF. In the volcano plots (Fig. 2j, 5d; Extended Data Fig. 3m), proteins were colored by class, all others were drawn in grey, and only significant proteins of a colored class were labelled. The principal component analyses (Extended Data Fig. 2n, 3l, 5e) used the Pol II, Integrator, initiation and elongation classes (105 gene symbols; 89 protein groups detected). Each comparison used the DMSO and treatment samples (5+5) separately, restricted to proteins quantified in all ten samples (86, 85 and 84 for INTS11-dTAG, INTS11-dTAG+NVP2 and NVP2). Median-normalized log2 intensities were centered and scaled to unit variance per protein before prcomp (R stats), and ellipses are 95% normal-theory data ellipses per condition (stat_ellipse, type = ‘norm’). The Integrator-subunit panel (Extended Data Fig. 2o) shows the limma log2 fold change (INTS11-dTAG versus DMSO) of the 15 detected subunits, with filled symbols marking adjusted P < 0.05 and no fold-change cutoff.

### Annotation

For all sequencing analyses, the human genome and annotation were obtained from UCSC: the hg38 (GRCh38) assembly, restricted to chromosomes 1-22, X, Y and M, and the NCBI RefSeq annotation (ncbiRefSeq; downloaded February 2024).

### Major isoform annotation

Major isoform annotation was performed with Salmon^92^ (v1.10.2). Salmon calculated transcript abundances in transcripts per million (TPM) by quantifying previously published HCT116 total RNA-seq data^93^ against the RefSeq transcriptome. Of the curated RefSeq transcripts, predicted models (XM, XR) and transcripts with TPM < 0.5 in both replicates were removed. The transcripts of each gene were grouped by exact TSS position. A gene was retained if its dominant TSS group carried more than 50% of the gene’s TPM, and the highest-TPM transcript of that group was taken as the major isoform. The final major isoform annotation excludes transcripts that overlap another major isoform on the same strand, except protein-coding hosts that fully contain a nested miRNA (13,220 expressed transcripts). Transcripts were further required to have at least 50 TT-seq^93^ read pairs per kilobase. This left 9,286 protein-coding and 846 other transcripts, including 21 snRNAs. Replication-dependent histone genes (45) were excluded from the mRNA class.

### uaRNAs and eRNAs

Upstream antisense RNAs (uaRNAs; n = 3,708) and eRNAs (n = 9,549) were taken from previous HCT116 TT-seq annotation^93^. uaRNAs were used without further filtering. eRNAs were removed if their body or 1 kb downstream flank overlapped an expressed major isoform (see above) on the same strand (8,816 remaining), or if they were shorter than 300 bp or longer than 5 kb (6,419 remaining). eRNAs were kept if their mean sense-strand TT-seq coverage in control cells (DMSO, 90 min; spike-in-normalized, mean of two replicates) was at least 0.3, which resulted in 3,861 eRNAs.

Enhancer DNA intervals were taken from previous HCT116 enhancer annotation^93^ (8,407 intervals). The 3,638 intervals that overlapped at least one of the 3,861 eRNAs on either strand were kept as active enhancers (median width 1,250 bp). Super-enhancers (SE) were the 49 regions defined in the MCC data (see MCC analysis). eRNAs that overlapped an SE region on either strand were classed as SE (n = 102). Typical-enhancer (TE) eRNAs were defined as the remaining eRNAs, excluding, in addition to the MCC-defined SEs, eRNAs located within HCT116 super-enhancers annotated in SEdb 3.0^94^ (SE_02_0352), which resulted in n = 3,140 TE eRNAs. An active-enhancer interval was classed as SE if any of its eRNAs was SE (n = 95 SE and n = 3,543 TE intervals).

### Metagene profiles and heatmaps

Profiles were computed from the merged normalized tracks (sense-strand for the stranded RNA assays, unstranded otherwise), anchored at the TSS of mRNAs or eRNAs or at peak centers (±1.5 kb), or, for TT-seq gene bodies, scaled into 600 bins between the TSS and the pA site/TES with 200-bin flanks. Each position shows the 5% trimmed mean across features (untrimmed for Pol II at eRNA TSSs), and the band is the 95% bootstrap confidence interval of that mean over features (500 resamples; boot package), so it reflects spread across features. Features whose window ran past a chromosome end were dropped. Readthrough profiles kept transcripts of at least 1 kb (mRNAs) or 200 bp (uaRNA, eRNA) with a complete, same-strand-free downstream window (20 kb or 5 kb) and were each normalized to the condition’s own signal just upstream of the pA site or TES. Heatmaps show the same matrices (±1.5 kb, 6 bp bins), rows sorted by the control signal over the central 10% of the window, with the color scale capped per panel pair at the higher of the two conditions’ 99th percentiles.

### Differential analysis and MA plots

For TT-seq and chrRNA-seq, read pairs were counted per RNA class (mRNA, snRNA, uaRNA, eRNA) with summarizeOverlaps (mode Union, strand-aware, duplicates excluded), each class in a separate pass. All conditions of a dataset were fitted together in DESeq2 (design ∼ replicate + condition, size factors fixed to the spike-in values); features with at least 5 reads in at least 2 samples were kept, contrasts taken by Wald test, log2 fold changes shrunk with ashr, and P values Benjamini-Hochberg-adjusted. Pol II core-promoter MA plots were computed from fragments shorter than 70 bp over the TSS −50 to +50 bp window of mRNAs, uaRNAs and eRNAs in a single DESeq2 fit (∼ condition) with Drosophila-based size factors and without shrinkage.

### Box plots and further tests

Boxes show the per-feature shrunken DESeq2 log2 fold changes; comparisons across cell lines or datasets were restricted to features passing the count filter in every dataset shown, and P values were adjusted within each panel. Pol II body fold changes (Extended Data Fig. 3n) used the mean Drosophila-normalized coverage of fragments shorter than 70 bp over the transcribed body, excluding the first 200 bp (mRNA, uaRNA) or 50 bp (eRNA) to remove the pause peak, with a half-minimum pseudocount.

### Elongation-velocity proxy

For Extended Data Fig. 4a, an elongation-velocity proxy was calculated for 3,283 eRNAs of at least 500 bp, excluding the first 100 bp from the TSS. TT-seq and Pol II xChIP-seq body coverages with assay-specific spike-in normalization were used. Per-assay pseudocounts were used as the median non-zero control coverage, and the proxy was log2[(TT-seq + pTT)/(Pol II xChIP-seq + pPolII)]. INTS11-dTAG and INTS11-dTAG+NVP2 were compared using a paired two-sided Wilcoxon signed-rank test, with Benjamini-Hochberg correction within each panel.

### Pol II pausing index

Computed per protein-coding transcript from Drosophila-normalized coverage of fragments shorter than 70 bp, as the mean in the promoter-proximal window (TSS +20 to +80 bp) over the mean in the body (TSS +300 bp to TES −300 bp). Transcripts shorter than 800 bp and replication-dependent histones were excluded. Transcripts were retained if their mean coverage was at least 0.5 in both windows in the control (DMSO). A pseudocount equal to the median non-zero control body coverage was added to both numerator and denominator before calculating the log2 pausing index.

### Public data analysis

DNase I hypersensitivity data^95^ and ChIP-seq data for H3K27ac^96^ and CTCF^95^ in HCT116 cells are available from ENCODE under accession codes ENCSR000ENM, ENCSR661KMA, and ENCSR000BSE, respectively, and ChIP-seq data for MED26^97^ are available from GEO under accession code GSE121355. Previously published HCT116 eRNA, uaRNA and enhancer annotations and HCT116 TT-seq and total RNA-seq data^93^, which were used to build the transcript and enhancer annotations (see Major isoform annotation and uaRNAs and eRNAs sections), are available from GEO under accession code GSE152291. HCT116 RAD21 ChIP-seq IDR-thresholded peaks, which were used to restrict the RAD21 ChIPmentation peak set, are available from ENCODE under accession code ENCFF391AAM. The ENCODE GRCh38 blacklist is available via accession number ENCFF356LFX. Structures of PIC-core Mediator and cohesin are available from PDB under PDB 7NVR and PDB 6WG3, respectively.

## ACKNOWLEDGEMENTS

We would like to thank Patrick Cramer for infrastructure support; Steven West for sharing the HCT116 INTS11-dTAG and HCT116 WDR82-mAID cell lines; Ibrahim Cissé, together with Oliver Siontas, Dimitra Tsouraki, Haneul Yoo, for advice on imaging experiments; the Facility for Mass Spectrometry and the Facility for Light Microscopy at the Max Planck Institute for Multidisciplinary Sciences, as well as the Flow Cytometry & DNA Sequencing Facility at the Planck Institute of Immunobiology and Epigenetics, for experimental support; Arjun Devadas and Yi Zhu for advice on bioinformatic analysis; Birgit Koschorz, Kerstin Maier, Ute Neef, Ralf Pflanz, Monika Raabe, Myriam Rohm, Petra Rus, Nicholas Shukeir, Lea Siegmund, and Robin Walz for experimental support; Argyris Papantonis and Johannes Söding for helpful feedback on the manuscript; and all members of the Oudelaar and Cramer departments for helpful discussions and feedback. This work was supported by the Max Planck Society; the European Research Council (Starter Grant 3D-REG 101115401); the Deutsche Forschungsgemeinschaft (DFG) via SFB 1565 (project 469281184/P02); the MSc/PhD program “Molecular Biology”— International Max Planck Research School at the Georg August University Göttingen; and the PhD program “Genome Science”—International Max Planck Research School at the Georg August University Göttingen.

## AUTHOR CONTRIBUTIONS STATEMENT

S.S. generated cell lines; performed MCC, Tiled-MCC, ATAC-seq, ChIPmentation and imaging experiments; analyzed data; prepared figures; and wrote the manuscript. T.V. conceived the project; performed TT-seq, chrRNA-seq, xChIP-seq and proteomics experiments; analyzed data; prepared figures; and wrote the manuscript. L.L.C. and S.S. contributed to imaging experiments and imaging data analysis. J.N. and J.L.W. performed biochemical reconstitution experiments. Y.Y., O.D. and H.U. performed proteomics experiments and analyzed proteomics data. S.R. and K.Z. provided conceptual advice. L.T.D and F.B. contributed to imaging experiments. M.L. conceived and supervised the project and wrote the manuscript. A.M.O. conceived and supervised the project; acquired funding; and wrote the first draft of the manuscript. All authors provided feedback on the manuscript.

## EXTENDED DATA FIGURES

**Extended Data Fig. 1.**
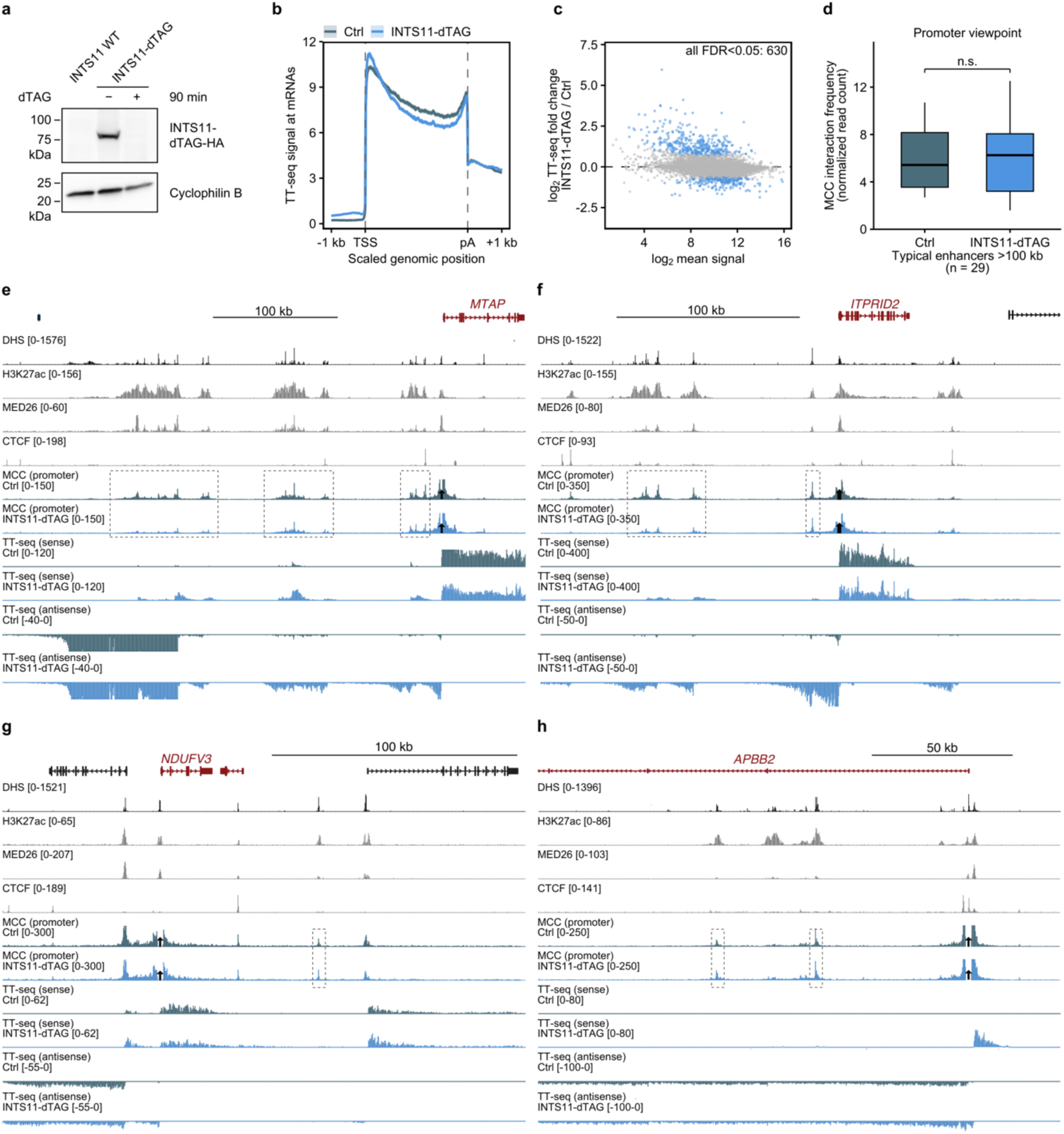
INTS11 depletion, protein-coding transcription and enhancer-promoter interactions at further loci. **a,** Western blot of HA-tagged INTS11-dTAG (anti-HA) in parental (WT) and INTS11-dTAG cells upon DMSO (-) or dTAG (+) treatment for 90 min; Cyclophilin B, loading control. **b**, TT-seq signal across protein-coding genes (n = 9,283), scaled between the TSS and the polyadenylation (pA) site, in Ctrl (grey) and INTS11-dTAG (blue). Profiles as in Fig. 1b. **c**, MA plot of TT-seq changes in protein-coding genes (n = 9,216) in INTS11-dTAG. Blue, adjusted P < 0.05; the number indicates the number of genes with adjusted P < 0.05. **d**, MCC interaction frequencies between promoter viewpoints and typical enhancers located >100 kb from the promoter (n = 29) in Ctrl and INTS11-dTAG. Box plots and tests as in Fig. 1g. P = 0.93; n.s., not significant. **e-h**, MCC interaction profiles (mean of n = 3 biological replicates) from the promoter viewpoints of MTAP (**e**; hg38 chr9:21,452,083-21,869,673) and ITPRID2 (**f**; chr2:181,726,563-182,014,405), genes with super-enhancers, and of NDUFV3 (**g**; chr21:42,831,184-43,037,379) and APBB2 (**h**; chr4:41,061,688-41,247,203), genes with typical enhancers in Ctrl (grey) and INTS11-dTAG (blue), with TT-seq, DNase I hypersensitive sites (DHS), H3K27ac, MED26 and CTCF.

**Extended Data Fig. 2.**
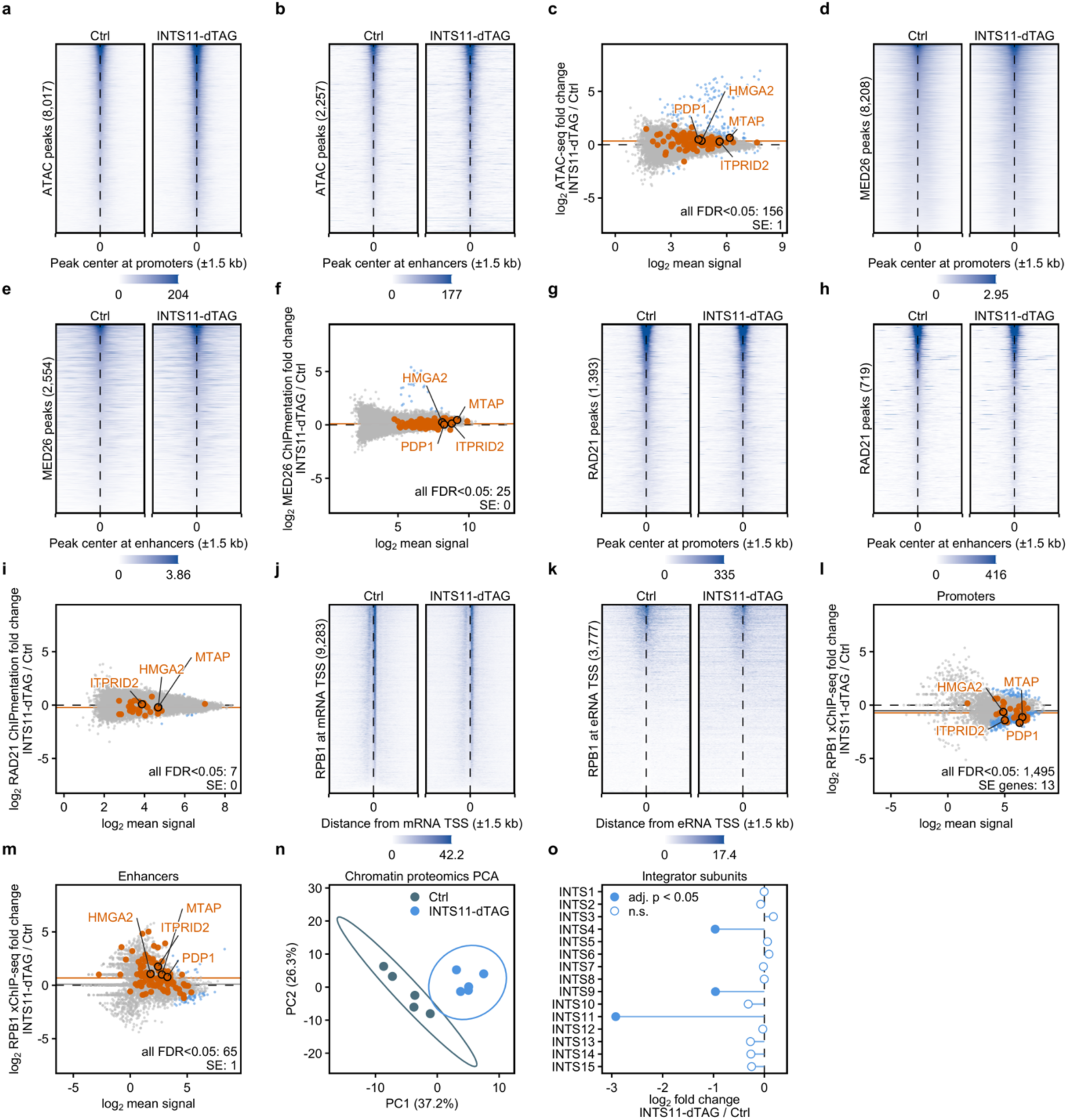
Chromatin accessibility, Mediator, cohesin and Pol II after INTS11 depletion. **a, b, d, e, g, h, j, k**, Heatmaps in Ctrl (left) and INTS11-dTAG (right) of ATAC-seq at promoters (**a**; n = 8,017) and enhancers (**b**; n = 2,257), MED26 ChIPmentation at promoters (**d**; n = 8,208) and enhancers (**e**; n = 2,554), RAD21 ChIPmentation at promoters (**g**; n = 1,393) and enhancers (**h**; n = 719), and RPB1 xChIP-seq at mRNA (**j**; n = 9,283) and eRNA (**k**; n = 3,777) TSSs, sorted by the control signal. **c, f, i, l, m**, MA plots of ATAC-seq (**c**; n = 47,077 peaks; 103 in MCC-targeted super-enhancers (SEs)), MED26 (**f**; n = 57,823; 152 in MCC-targeted SEs), RAD21 (**i**; n = 39,514; 31 in MCC-targeted SEs), and Pol II in the core promoter (TSS ± 50 bp) of mRNAs (**l**; n = 9,268; 30 MCC-targeted SE-associated genes) and eRNAs (**m**; n = 3,679; 102 MCC-targeted SE eRNAs) in INTS11-dTAG. Blue, adjusted P < 0.05; orange, SE features in MCC-targeted loci (peaks overlapping an SE enhancer or SE eRNA; in **l**, the SE-associated genes) and their median. The numbers give the features with adjusted P < 0.05 among all features and among MCC-targeted SE features. For HMGA2, PDP1, MTAP and ITPRID2, the SE feature with the highest mean signal is labelled (black ring). MA plots here and in Extended Data Fig. 3 are plotted on the same y-axis to allow for direct comparison. **n**, Principal component analysis of Pol II-associated proteins (n = 86) in Ctrl (grey) and INTS11-dTAG (blue); ellipses are 95% normal data ellipses; n = 5 biological replicates. **o**, Chromatin-binding changes of Integrator subunits in INTS11-dTAG. Filled dots, adjusted P < 0.05.

**Extended Data Fig. 3.**
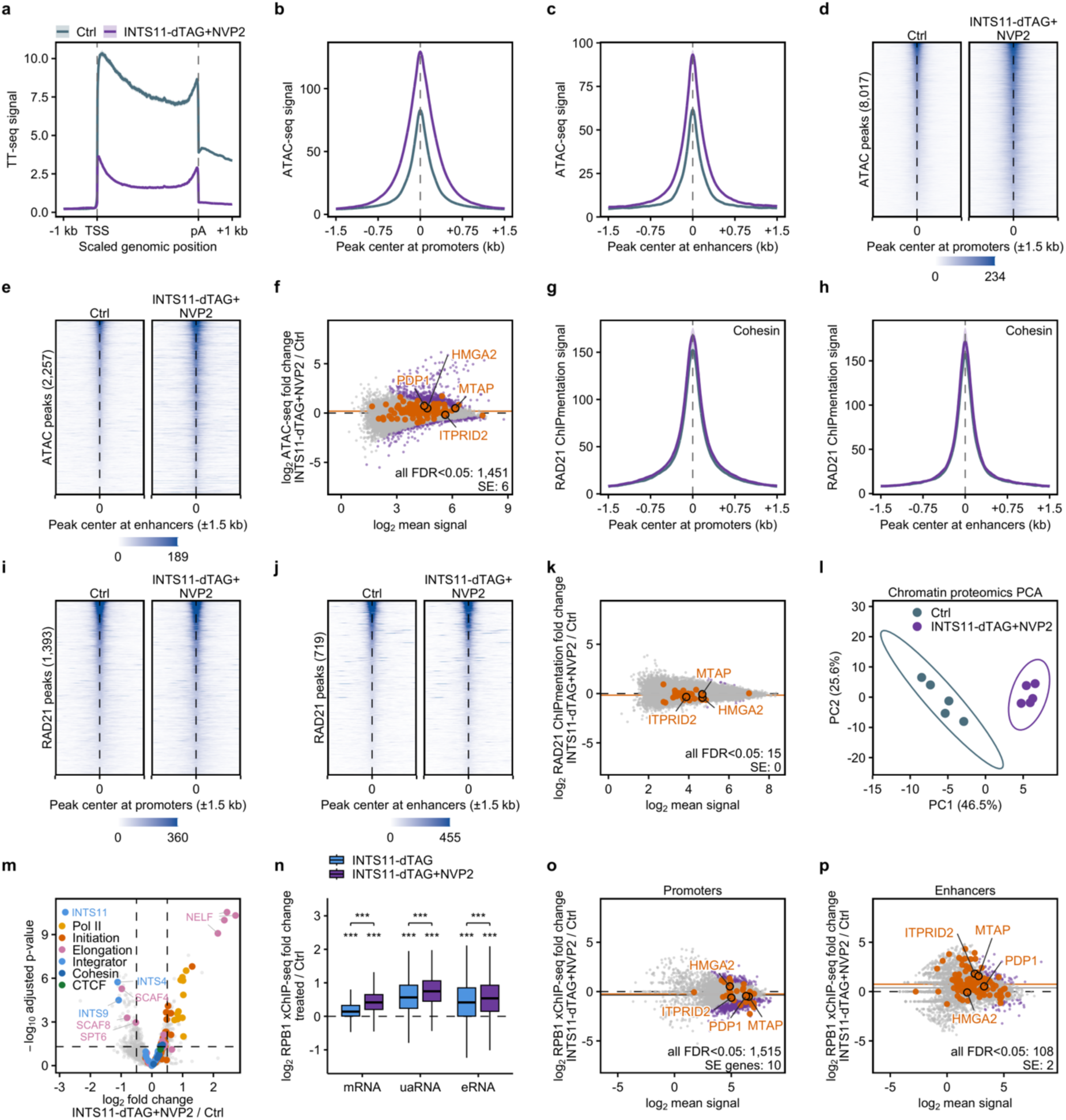
CDK9 inhibition in INTS11-depleted cells. All panels compare Ctrl (grey) with INTS11-dTAG+NVP2 (purple), except n. **a**, TT-seq signal across protein-coding genes (n = 9,283), scaled between the TSS and the pA site. Profiles as in Fig. 1b. **b, c**, ATAC-seq at promoters (**b**; n = 8,017) and enhancers (**c**; n = 2,257), as in Extended Data Fig. 2. **d, e**, ATAC-seq heatmaps, as in Extended Data Fig. 2. **f**, ATAC-seq MA plot (n = 47,077 peaks; 103 in MCC-targeted SEs), as in Extended Data Fig. 2. Purple, adjusted P < 0.05. **g, h**, RAD21 ChIPmentation at promoters (**g**; n = 1,393) and enhancers (**h**; n = 719), as in Extended Data Fig. 2i. **i, j**, RAD21 heatmaps, as in Extended Data Fig. 2. **k**, RAD21 MA plot (n = 39,514 peaks; 31 in MCC-targeted SEs), as in Extended Data Fig. 2. **l**, Principal component analysis of Pol II-associated proteins (n = 85). **m**, Chromatin proteomics volcano plot (4,454 proteins), as in Fig. 2j; 94 proteins were enriched on and 132 lost from chromatin. **n**, RPB1 xChIP-seq fold changes in the bodies of mRNAs (n = 9,280), uaRNAs (n = 3,422) and eRNAs (n = 3,860) in INTS11-dTAG (blue) or INTS11-dTAG+NVP2 (purple). Box plots and tests as in Fig. 3a. **o, p**, RPB1 core-promoter MA plots of mRNAs (**o**; n = 9,268) and eRNAs (**p**; n = 3,679), as in Extended Data Fig. 2.

**Extended Data Fig. 4.**
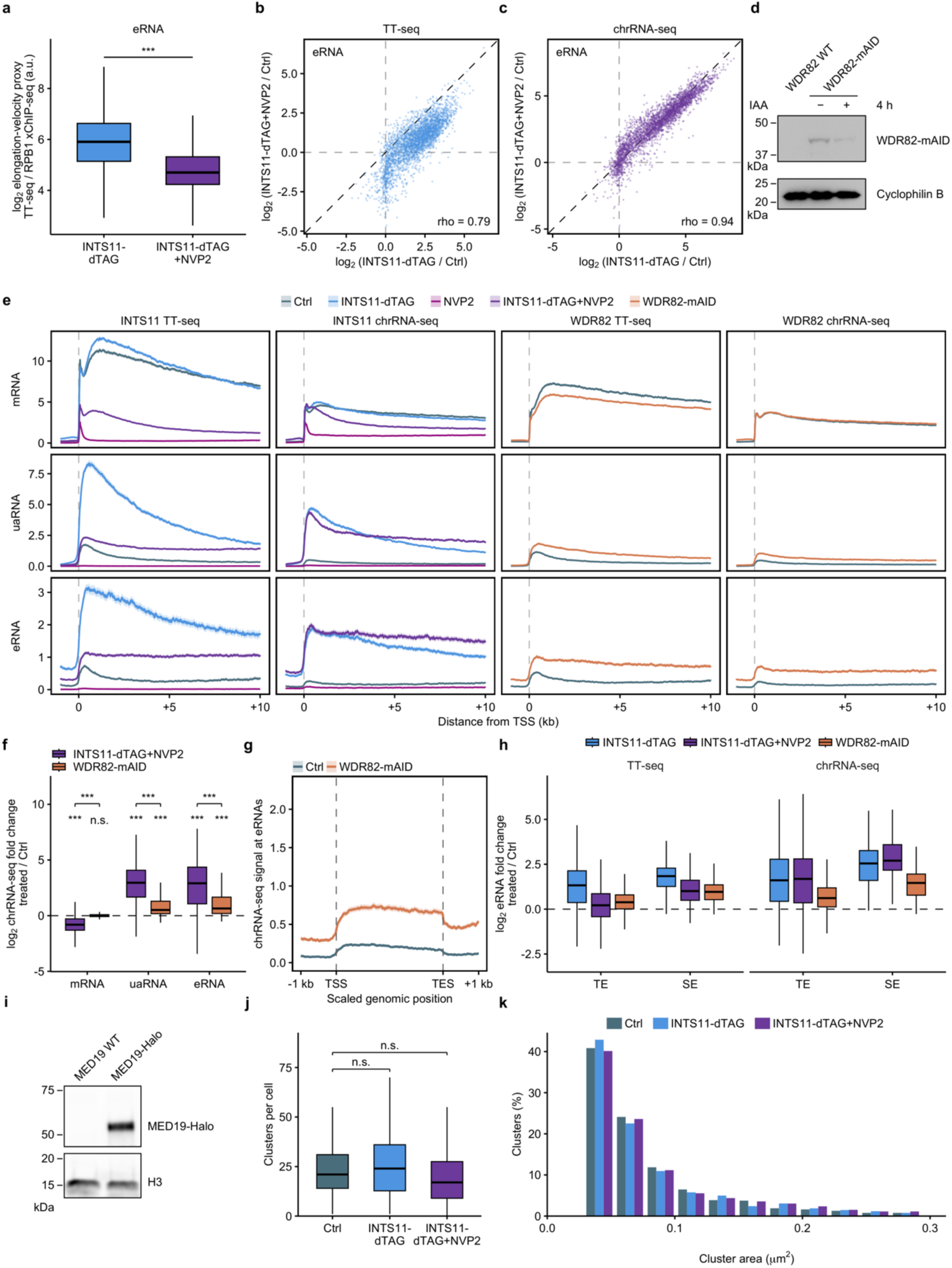
Comparison of INTS11 and WDR82 perturbations. **a,** eRNA elongation-velocity proxy, log2(TT-seq / RPB1 xChIP-seq), in INTS11-dTAG and INTS11-dTAG+NVP2 (n = 3,283 eRNAs). Two-sided Wilcoxon signed-rank test, paired per eRNA. Box plots as in Fig. 1g. **b, c**, eRNA fold changes (n = 3,076) in INTS11-dTAG (x-axis) and INTS11-dTAG+NVP2 (y-axis) as measured by TT-seq (**b**) and chrRNA-seq (**c**), with Spearman correlation coefficients; dashed line indicates equal response. **d**, Western blot of WDR82-mAID (anti-mAID) in parental (WT) and WDR82-mAID cells after DMSO (-) or IAA (+) treatment for 4 h; Cyclophilin B, loading control. **e**, TT-seq and chrRNA-seq signal at the TSSs of mRNAs (top; n = 9,283), uaRNAs (middle; n = 3,706) and eRNAs (bottom; n = 3,860) for INTS11-dTAG (blue), NVP2 (magenta), INTS11-dTAG+NVP2 (purple) and WDR82-mAID (orange), each with its own control (grey). Profiles as in Fig. 1b. **f**, chrRNA-seq fold changes in INTS11-dTAG+NVP2 (purple) or WDR82-mAID (orange), each against its own control, of mRNAs (n = 9,189), uaRNAs (n = 2,268) and eRNAs (n = 1,831) detected in both conditions. Box plots and tests as in Fig. 3a. **g**, chrRNA-seq signal across eRNAs (n = 3,860) in Ctrl (grey) and WDR82-mAID (orange). Profiles as in Fig. 4b, plotted on the same y-axis to allow for direct comparison. **h**, TT-seq (left) and chrRNA-seq (right) fold changes of eRNAs at typical enhancers (TE; n = 3,140) and MCC-targeted super-enhancers (SE; n = 102) in INTS11-dTAG, INTS11-dTAG+NVP2 or WDR82-mAID. Box plots as in Fig. 1g. **i**, Western blot of HaloTag-tagged MED19 (anti-Halo) in INTS11-dTAG cells with wild-type MED19 (MED19 WT) or the MED19-Halo knock-in (MED19-Halo); H3, loading control. **j**, SIM MED19-HaloTag clusters per cell in Ctrl, INTS11-dTAG and INTS11-dTAG+NVP2 (n = 94, 92 and 111 cells; clusters of 30-500 pixels, 0.029-0.48 µm²). Two-sided Wilcoxon rank-sum tests against Ctrl, Benjamini-Hochberg adjusted; P = 0.50 and 0.15. Box plots as in Fig. 1g. **k**, SIM MED19-HaloTag cluster size distribution expressed as percentage per 25-pixel (0.024 µm^2^) area bin; axis cut at 0.3 µm^2^ (1.8-2.4% of clusters lie beyond). n.s., not significant; ***P < 0.001.

**Extended Data Fig. 5.**
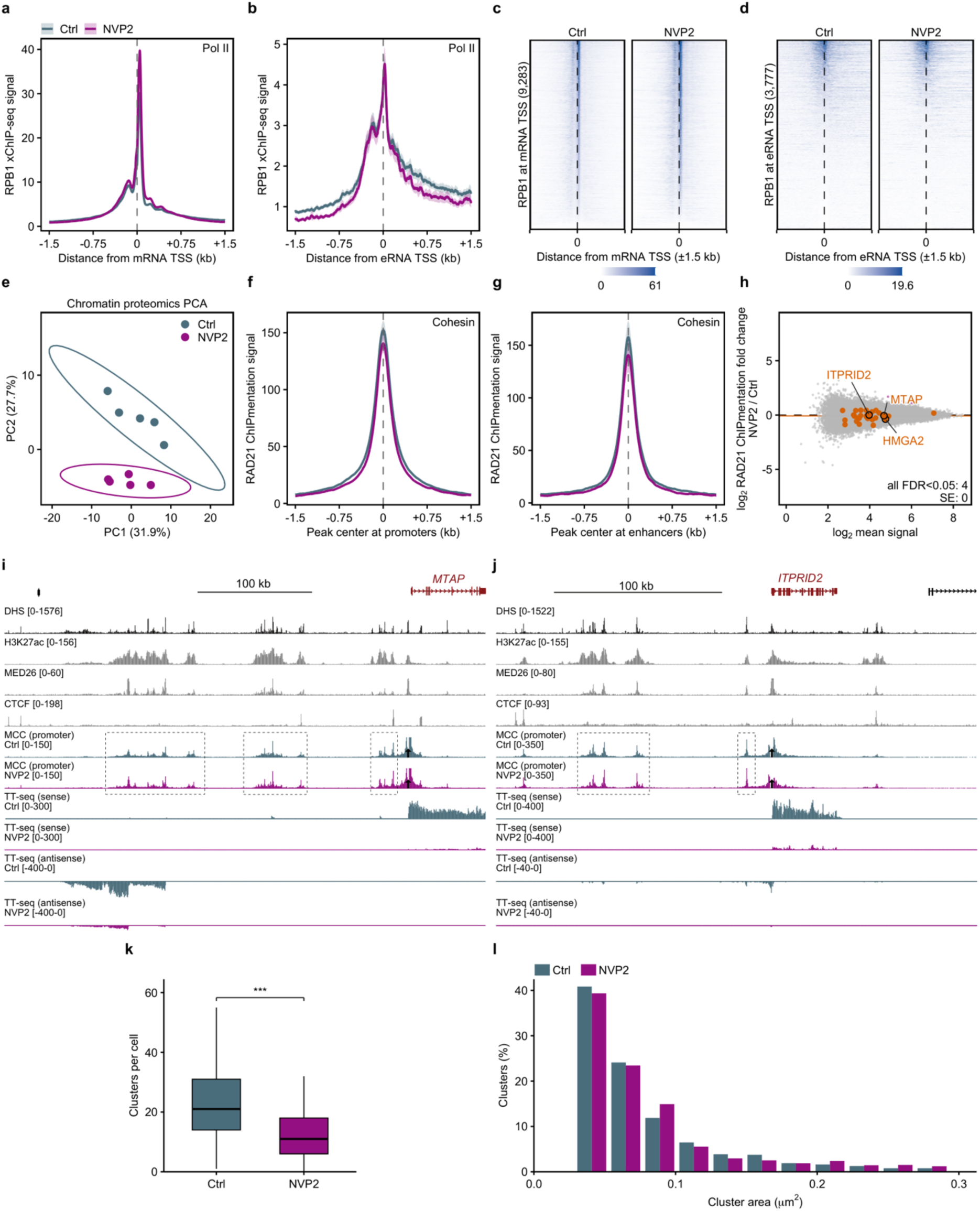
Pol II, cohesin and enhancer-promoter interactions upon CDK9 inhibition. All panels compare Ctrl (grey) with NVP2 for 45 min (magenta). **a, b**, RPB1 xChIP-seq signal at mRNA (**a**; n = 9,283) and eRNA (**b**; n = 3,777) TSSs, as in Fig. 2h,i. **c, d**, RPB1 heatmaps, as in Extended Data Fig. 2. **e**, Principal component analysis of Pol II-associated proteins (n = 84), as in Extended Data Fig. 2n. **f, g**, RAD21 ChIPmentation at promoters (**f**; n = 1,393) and enhancers (**g**; n = 719), as in Fig. 2f,g. **h**, RAD21 MA plot (n = 39,514 peaks; 31 in SEs), as in Extended Data Fig. 2; magenta, adjusted P < 0.05. i, j, MCC interaction profiles from the promoter viewpoints of MTAP (**i**; hg38 chr9:21,452,083-21,869,673) and ITPRID2 (**j**; chr2:181,726,563-182,014,405), with TT-seq signal; n = 3 MCC biological replicates per condition. Tracks as in Fig. 1e. **k**, SIM MED19-HaloTag clusters per cell in Ctrl (n = 94 cells) and NVP2 (n = 104 cells), as in Extended Data Fig. 4j. P = 8.7 × 10^−9^. **l**, SIM MED19-HaloTag cluster size distribution, as in Extended Data Fig. 4k; 2.4-2.7% of clusters lie beyond the axis. ***P < 0.001.

**Extended Data Fig. 6.**
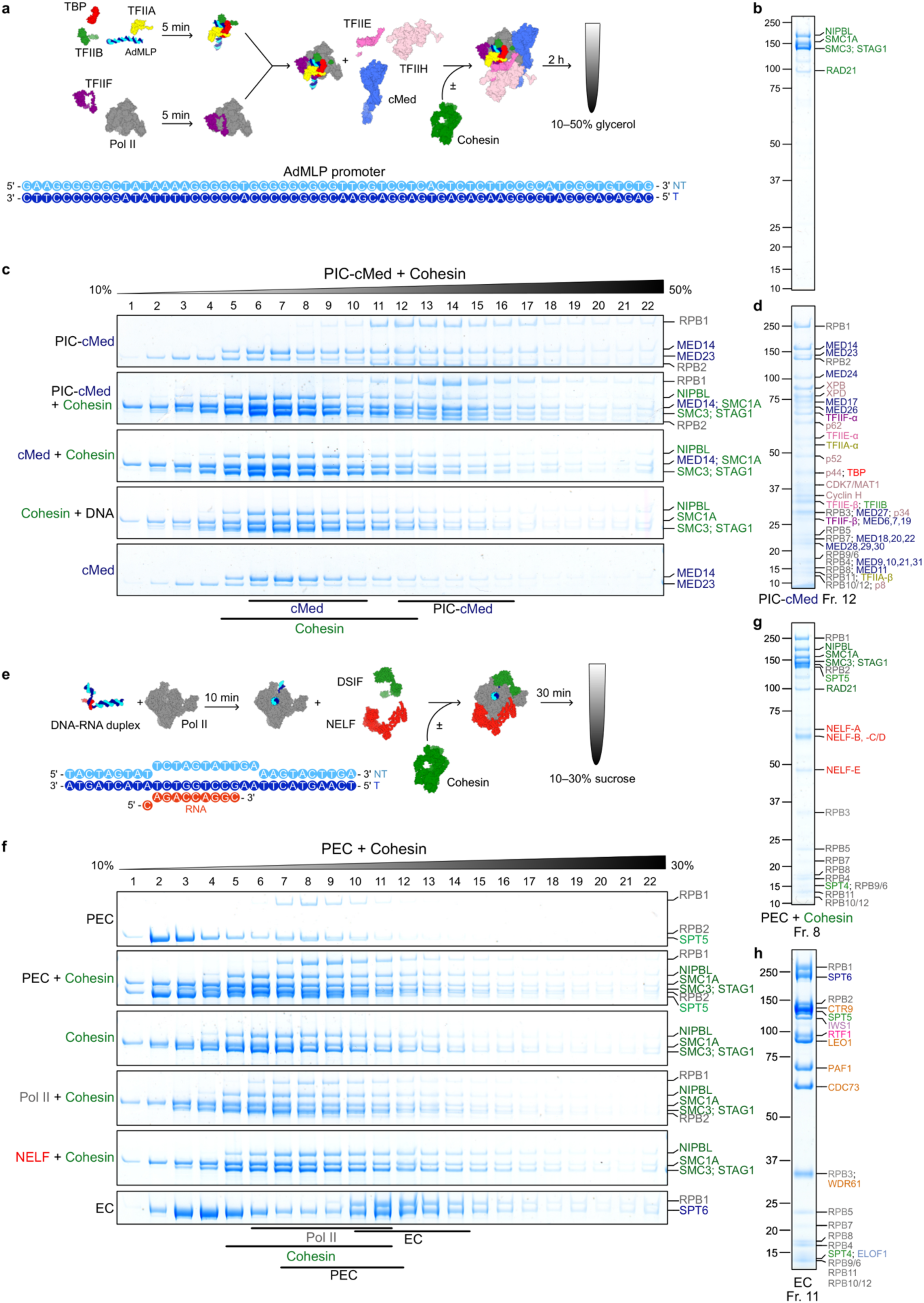
Cohesin does not form stable complexes with the PIC-Mediator complex or the paused elongation complex in vitro. **a,** Assembly of the Pol II pre-initiation complex (PIC) with core Mediator (cMed; together PIC-cMed) on the TATA-box-containing adenovirus major late promoter (AdMLP), with and without cohesin, before density gradient ultracentrifugation; structures from PDB 7NVR (PIC-cMed) and 6WG3 (cohesin-NIPBL). **b**, SDS-PAGE of purified cohesin (SMC1A, SMC3, RAD21 and STAG1) in complex with NIPBL. **c**, Glycerol gradients (10-50%) of the indicated PIC-cMed compositions with and without cohesin; peak fractions of cohesin, cMed, and PIC-cMed are indicated. All fractions were analyzed by SDS-PAGE. A stable PIC-cMed-cohesin complex would shift cohesin into heavier fractions and co-sediment with PIC-cMed. However, cohesin peaks in the same fraction with and without PIC-cMed, indicating that no stable complex forms. **d**, Example of a single gradient fraction: SDS-PAGE of peak fraction 12 of the PIC-cMed gradient in **c**, showing all expected components. **e**, Assembly of the paused elongation complex (PEC; Pol II with DSIF and NELF) on a DNA-RNA scaffold, with and without cohesin, before density gradient ultracentrifugation; structure from PDB 6GML. **f**, Sucrose gradients (10-30%) of Pol II and PEC compositions with and without cohesin; peak fractions of Pol II, cohesin, and PEC are indicated. An activated elongation complex (EC; Pol II with DSIF, SPT6, IWS1, RTF1, PAF1 complex, ELOF1 and P-TEFb; 1.3 MDa) is the size reference; a PEC-cohesin complex would be 1.4 MDa and would therefore sediment beyond the EC peak. **g**, As in **d**, SDS-PAGE of fraction 8 of the PEC-cohesin gradient in f, which contains both PEC and cohesin because the two sediment in overlapping fractions. No species heavier than the PEC is observed at the position expected for a PEC-cohesin complex, indicating that no stable complex forms. **h**, As in **d**, SDS-PAGE of peak fraction 11 of the EC gradient in f.

